# Characterizing the Ion-Conductive State of the *α*7-Nicotinic Acetylcholine Receptor via Single-Channel Measurements and Molecular Dynamics Simulations

**DOI:** 10.1101/2025.08.15.670429

**Authors:** Nauman Sultan, Gisela D. Cymes, Ada Chen, Bernard Brooks, Claudio Grosman, Ana Damjanovic

**Affiliations:** Department of Physics & Astronomy, Johns Hopkins University, Baltimore, MD 21218, U.S.A.; National Heart, Lung, and Blood Institute; National Institutes of Health, Bethesda, MD 20892, U.S.A.; Department of Molecular and Integrative Physiology, University of Illinois Urbana-Champaign, Urbana, IL 61801, U.S.A.; Center for Biophysics and Quantitative Biology, University of Illinois Urbana-Champaign, Urbana, IL 61801, U.S.A.; Neuroscience Program, University of Illinois Urbana-Champaign, Urbana, IL 61801, U.S.A.; Department of Biophysics, Johns Hopkins University, 3400 N. Charles Street, Baltimore, MD 21218, U.S.A.

## Abstract

The α7-nicotinic acetylcholine receptor (α7-nAChR) is a cation-selective Cys-loop receptor involved in diverse physiological processes and is an important therapeutic target. Multiple cryo-EM structures of putative open states are now available and their functional relevance is under active investigation. Here, we combined single-channel patch-clamp recordings with atomistic molecular dynamics (MD) simulations to assess the conductive properties of several α7-nAChR structures solved with different ligands. Simulations restrained to the respective cryo-EM structures produced only modest ion flux for all models, inconsistent with experiment, whereas fully unrestrained simulations revealed marked differences in their ability to relax into physiologically conductive ensembles. Two structures, 7KOX and ligand-bound 8V82, consistently stabilized into conductive states whose permeation properties matched our measured inward single-channel conductance. The conduction of 8V82 nearly stopped upon removing the modeled ligands. 8V80 showed only intermittent conduction with ligands and remained non-conductive without them. 7EKT collapsed into a non-conductive conformation upon relaxation, irrespective of whether the modeled ligands were retained or removed. 9LH5, despite having a transmembrane pore nearly identical to 7KOX’s, exhibited approximately twofold higher conductance, likely due to a widened extracellular vestibule. Across models, permeation events followed Poissonian statistics with a characteristic entry lag captured by a double-Poisson model. Simulations of outward currents consistently overestimated the conductance compared to experiments, perhaps reflecting the absence of the full intracellular domain in available structural models and/or the presence of current-blocking concentrations of cytosolic Mg^2+^ in patch-clamp cell-attached recordings. These results identify the conformations most compatible with the physiological open state and underscore the importance of unrestrained MD, ligand stabilization, and extracellular-vestibule geometry in shaping α7-nAChR conduction.

## Introduction

The nicotinic acetylcholine receptors (nAChRs) are neurotransmitter-gated ion channels that transduce the binding of extracellular acetylcholine (ACh) into the passive flow of inorganic cations into the cell. Along with homologous receptors to serotonin, γ-aminobutyric acid, glycine, and Zn^2+^, the nAChRs constitute the superfamily of Cys-loop receptors of vertebrate animals^1–3^. These membrane receptors are pentameric assemblies of identical or homologous subunits arranged around a central pore. The extracellular domain (ECD) harbours the neurotransmitter-binding sites, the transmembrane domain (TMD) forms the transmembrane pore, and five short stretches of amino acids—one per subunit—link the two domains covalently^4^ ensuring their functional coupling^5^.

In vertebrates, nAChRs are formed by a variety of subunits (α1–α10, β1–β4, γ, δ, and ε), α7 being one of the few that can assemble into fully functional homomers. Another unique aspect of the α7-nAChR is its expression in many different cell types of the animal body, which underlies its participation in a wide range of physiological phenomena and pathological conditions.

All neurotransmitter-gated ion channels (whether from the superfamily of Cys-loop receptors or not) can interconvert among a variety of conformations that differ in their ability to conduct ions and the affinity with which they bind neurotransmitters^6^. The conformational space accessible to these receptor-channels is typically complex, but it minimally consists of three end states denoted as “closed”, “open” and “desensitized”. The open state conducts ions, whereas the closed and desensitized states do not. On the other hand, the open and desensitized states bind neurotransmitter with a higher affinity than the closed state does ^6^. In the absence of neurotransmitter, the closed state is the most stable conformation, and its occupancy approaches unity. However, because of their different neurotransmitter-binding affinities, the presence of neurotransmitter favours the open and desensitized states. Indeed, in the presence of saturating concentrations of neurotransmitter, the equilibrium occupancy of the closed state drops to nearly zero. Since the neurotransmitter-bound desensitized state is much more stable than the neurotransmitter-bound open state, the occupancy of the latter at equilibrium is, also, typically very low. As a result, the neurotransmitter-bound open state is only transiently occupied as the concentration of neurotransmitter is stepped from zero to saturating. It follows then that, in the absence of neurotransmitter, nearly all channels are closed, whereas in the presence of saturating concentrations of neurotransmitter, nearly all channels are desensitized.

In recent years, largely through the application of cryogenic electron microscopy (cryo-EM), the structures of all major types of Cys-loop receptors in at least one conformation have been solved. Not surprisingly, the unliganded closed state is the conformation most commonly isolated in the absence of neurotransmitter (or any other efficacious agonist that binds to the neurotransmitter-binding sites), and the agonist-bound desensitized state is the conformation most commonly observed in the presence of saturating full agonists. Different approaches have been followed to enrich the mixture of conformational states in the elusive agonist-bound open conformation, with the millisecond time-scale, out-of-equilibrium method of Unwin and coworkers (“plunge-freezing” ^7^) having proved to be the very successful^8,9^. However, a few atomic models of Cys-loop receptors obtained under equilibrium conditions have been proposed to exhibit features consistent with agonist-bound open conformations; such is the case for the α7-nAChR, for example. A plausible explanation for this type of finding is that the lipid-membrane mimetic used to maintain the protein in solution for cryo-EM imaging may alter the channel’s conformational free-energy landscape relative to that in a native plasma membrane. Whatever the underlying reason might be, an atomic model of the receptor’s open state affords the opportunity of studying the phenomenon of biological ion conduction using atomistic molecular dynamics (MD) simulations.

Here, we present simulations and an electrophysiological characterization of the open-channel state of the human α7-nAChR. Since Na^+^-ion permeation through the α7-nAChR has been characterized in previous studies^10–13^, our analysis for this cation is focused on demonstrating close agreement between the computed and experimentally measured single-channel conductances. Instead, here we focused on the conduction of K^+^ ions to provide a complementary view. K^+^ ions also diffuse faster than Na^+^-ions in water, facilitating better overall statistics and identification of rare pathways (e.g., lateral fenestration) in finite-time simulations. We performed all-atom MD simulations to gain mechanistic insight into the phenomenon of K^+^-ion permeation through five atomic models of this receptor that were deemed to represent open or open-like states: PDB ID 7KOX, ^14^ 7EKT,^15^ 8V80,^16^ 8V82,^16^ and 9LH5.^17^ The first four structures were obtained under conditions that slow down (but, certainly, do not prevent) desensitization, that is, in the presence of both orthosteric agonists and TMD-binding modulators. 9LH5, on the other hand, was obtained in the presence of a sub-saturating concentration of orthosteric agonist, only. We ran MD simulations of all the atoms resolved in the extracellular, transmembrane and intracellular parts of the channel. The intracellular region of the α7-nAChR^18^ was not fully modeled in any of these PDBs and consequently was absent from the MD setup. Simulations were restrained to retain the published coordinates ("PDB states") and two more independent sets of unrestrained simulations with and without bound ligands. From these, we computed the single-channel conductance values, tracking the pathway followed by the permeating ions. None of the restrained simulations produced single-channel conductances that matched the experimentally estimated value for the fully open state of the receptor. However, unrestrained simulations of two atomic models (PDB ID 7KOX and 8V82) did.

## Experimental and Computational Methods

### cDNA clones, Mutagenesis, and Heterologous Expression

Complementary DNA (cDNA) coding the human α7-nAChR (UniProt accession number: P36544) in pcDNA3.1 was purchased from addgene (#62276); cDNA coding isoform 1 of human RIC-3 (accession number: Q7Z5B4) ^19^ in pcDNA3.1 was provided by W. N. Green (University of Chicago, IL); cDNA coding human NACHO (TMEM35A; accession number: Q53FP2) ^20^ in pCMV6-XL5 was purchased from OriGene Technologies Inc. (#SC112910); and cDNAs coding the mouse α1, β1, δ, and ε subunits of the (muscle) nAChR (accession numbers: P04756, P09690, P02716, and P20782, respectively) in pRBG4 were provided by S. M. Sine (Mayo Clinic, Rochester, MN). The T264P mutation in the ε subunit^21^ (M2, position 12’) was engineered using the QuikChange kit (Agilent Technologies). Both the α7 and adult-muscle nAChRs were heterologously expressed in transiently transfected adherent HEK-293 cells grown at 37*^◦^*C and 5% CO_2_ in 35-mm cell-culture dishes. Transfections were performed using a calcium-phosphate method, and cells were incubated with the calcium-phosphate-DNA precipitate for 16-18 h after which the cell-culture medium (DMEM; Gibco) was replaced with fresh medium. For the expression of the human α7-nAChR, the transfection mixture contained a total of 3.0 µg/dish of cDNAs coding the α7-nAChR subunit, RIC-3, and NACHO in a 1:5.5:5.5 ratio (by weight). For the expression of the mouse adult-muscle nAChR, the transfection mixture contained a total of 0.75 micrograms/dish of cDNAs coding the α1, β1, δ, and ε subunits in a 2:1:1:1 ratio (by weight).

### Electrophysiology

Single-channel currents were recorded in the cell-attached configuration at ∼22*^◦^*C using an Axopatch 200B amplifier (Molecular Devices). The output signal was low-pass filtered (900C; Frequency Devices) so that the effective bandwidth was DC–30 kHz. Currents were digitized (at 100 kHz) and analyzed using QuB 1.4 software (the MLab Edition). For all recordings from the α7-nAChR„ the bath solution was 142 mM NaCl, 5.4 mM KCl, 1.8 mM CaCl_2_, 1.7 mM MgCl_2_, and 10 mM HEPES/NaOH, pH 7.4. For K^+^-current recordings, the pipette solution was 110 mM KCl, 40 mM KF, 1 or 100 µM ACh, 3 µM PNU-120596 (a positive allosteric modulator that slows down desensitization^22^), 0.1% v/v DMSO, and 10 mM HEPES/KOH, pH 7.4. For Na^+^-current recordings, the pipette solution was 150 mM NaCl, 100 µM ACh, 3 µM PNU-120596, 0.1% v/v DMSO, and 10 mM HEPES/NaOH, pH 7.4. For all recordings from the εT264P mutant of the muscle nAChR (a mutant with prolonged openings^21^), the bath solution was 142 mM KCl, 5.4 mM NaCl, 1.8 mM CaCl_2_, 1.7 mM MgCl_2_, and 10 mM HEPES/KOH, pH 7.4. For K^+^-current recordings, the pipette solution was 150 mM KCl, 1 µM ACh, and 10 mM HEPES/KOH, pH 7.4. For Na^+^-current recordings, the pipette solution was 150 mM NaCl, 1 µM ACh, and 10 mM HEPES/NaOH, pH 7.4. Patch pipettes, pulled from thick-walled borosilicate-glass capillary tubing (Sutter Instrument), had resistances of 7–10 MΩ when filled with pipette solution. Current amplitudes between the fully open and the zero-current (“shut”) levels were estimated from the idealization of selected stretches of single-channel activity using the segmental *K* -Means (SKM) method implemented in QuB^23^. These current amplitudes were used to generate current voltage plots, and the latter were used to estimate the single-channel conductance (as the slope of these linear plots). For display purposes, example single-channel traces were further low-pass filtered at 5 kHz.

### Molecular Dynamics Model

The starting points for our MD simulations were atomic models PDB IDs 7KOX, 7EKT, 8V80, 8V82, and 9LH5. All protein residues resolved in the structures were included in the simulation and are listed in the SI, Table S1. CHARMM-GUI^24–30^ was used to create the protein–membrane bilayer system and add ions. The generated systems were electrically neutral, featuring periodic boundary conditions in all three dimensions and a bulk ion concentration of 150 mM KCl or 150 mM NaCl. The protonation states of ionizable residues were modelled on the basis of CHARMM-GUI recommendations at pH 7, which use PROPKA^31,32^ calculations. In particular, all Asp and Glu residues were modelled as negatively charged, all Lys and Arg residues as positively charged, and all His residues were modeled as neutral. For 7KOX, the proton on His residues was kept on the Nitrogen Epsilon (NE) atom (7KOX PDB structure contained explicit hydrogen atom coordinates). For all other structures, the proton on His was modeled on the Nitrogen Delta (ND) atom, on the basis of recommendations by CHARMM-GUI. Glycans were not included in the MD simulations for any of the PDB models. The termini patching were standard N-(NTER) and C-terminus (CTER) for all models. The system setup details are included in SI, subsection S1.1.

As mentioned in the introduction, MD simulations of all PBD models were either restrained or unrestrained. For all restrained simulations, the modeled ligands were removed. Unrestrained simulations, on the other hand, were run with and without the modeled ligands. Ligands topology and parameter files were generated using CHARMM-GUI (using CHARMM General Force Field, CGenFF). Table 1 gives details of the PDB IDs and the ligands found in the structures. The membrane bilayer was composed of POPC lipids. Box dimensions, number of molecules, and disulfide-bond details are included in Table S1 to S3. The minimum distance between periodic images of protein atoms in *x* and *y* axes was at least ∼20 Å, and along the *z* -axis, it was ∼30 Å in every setup. During the setup, pore water was added up to a radius of 9 Å in all cases, and the channel was vertically aligned for membrane placement by running PPM 2.0^33^ in CHARMM-GUI. The five Ca^2+^ ions present in the PDB models 7KOX, 8V80, and 8V82 were restrained during the MD simulation to the C-α atoms of Glu 44 using a harmonic restraint.

**Table 1:** PDB IDs of the α7-nAChR used for the MD simulations, the lipid condition under which these were resolved, and the ligands bound to these models.

| PDB IDs | Lipid Mimetic | Orthosteric Ligand<br>(Bound to ECD) | Allosteric Modulator<br>(Bound to TMD) |
| --- | --- | --- | --- |
| 7EKT | Detergent Micelle | EVP-6124 | PNU-120596 |
| 7KOX | Nanodisc | Epibatidine | PNU-120596 (not resolved) |
| 8V80 | Detergent Micelle | Epibatidine | TQS |
| 8V82 | Detergent Micelle | Epibatidine | PNU-120596 |
| 9LH5 | Detergent Micelle | L-nicotine | None |

### Minimization and Equilibration

CHARMM36m^34^ was used as the force-field. The SHAKE algorithm^35^ was used to impose a holonomic restraint on bonds containing hydrogen atoms, and the SETTLE algorithm^36^ was used to constrain OH bonds in rigid water (TIP3P^37^) molecules.

Before the velocities were assigned to each atom, the energy of the system was minimized by using 5,000 steps of LocalEnergyMinimizer in OpenMM (the position was searched using the L-BFGS algorithm) or a combination of 2,500 steps of steepest descent and 2,500 steps of conjugate gradient in AMBER.

Particles in the minimized systems were assigned velocities from the Maxwell–Boltzmann distribution at a temperature of 30*^◦^*C (CHARMM-GUI default), and the systems were equilibrated. Equilibration proceeded as a 6-step process (for each replica in each setup), following the CHARMM-GUI protocol (see Table S2); at each step, the bond and dihedral restraints were gradually lifted until completely removed before the production run. In restrained simulations, protein atoms were restrained with a 10 kcal/mol/Å^2^ harmonic restraint (step 1 of the equilibration protocol) during the entire run.

Additional equilibration runs were performed for all unrestrained simulations started after running the system without restraints (see Table S4 to S6). This step was also carried out for restrained systems, but as mentioned before, the restraints on the atoms of the protein were kept fixed. The results and trajectories from minimization and equilibration steps were not included in the analysis. Two different ensembles were used to simulate the systems: NPT and NVT setups were used in OpenMM^38^, and only NVT was used in AMBER^39–42^. For NPT systems, the pressure was maintained at 100 kPa using a MonteCarlo (MC) barostat coupling. All systems were coupled to a heat bath using Langevin dynamics with friction coefficient of 1 ps to maintain the temperature at 30*^◦^*C (CHARMM-GUI default).

### Production Runs

Production simulations had an external electric field (EF) applied^43^, with strength adjusted to maintain the target membrane potential. The magnitude of the applied EF is included in Table S3. Each unrestrained production run was at least 200-ns long, with the longest run being 1 µs. The restrained simulations were all 500-ns long for each PDB; since the protein structure during these runs did not vary, only one copy was enough to obtain reliable statistics. Details regarding the length of production runs, ensembles, and programs used to simulate each setup can be found in Tables S4 to S8. The systems were integrated at a time step of 2 fs, and the coordinates of atoms were saved every 10 ps.

### Post Simulation Analysis

Visual Molecular Dynamics (VMD)^44^, along with repositories HOLE^45^ and PMEPot Plugin^46^ were used to calculate ion densities, channel radius, and the electrostatic-potential map of each system. CPPTRAJ^47,48^ was used to calculate position densities of residues, to combine trajectories, and to calculate average structures. Python code was written using MDTraj^49^ to calculate channel conductance, RMSDs between structures, and to plot ion trajectories.

#### Channel Conductance

Single-channel conductance was estimated by calculating the current at specified membrane potentials^46,50–52^. In turn, currents were calculated by counting the net number of charges crossing the membrane per unit time. The details of the method are included in SI, section S2.

#### Poisson Process to Define Events Distribution

If events (defined by a complete crossing of an ion through the TMD of the channel; see SI, section S2) are independent of one another in a disjoint time interval, and if the distribution of such events is defined by a single rate parameter *λ* for an interval *δt*, which can be translated for any time *T*, the distribution of events can be characterized by a homogeneous Poisson point process.

The probability density (PDF) and the cumulative distribution (CDF) functions for a Poisson process are, respectively, given by:

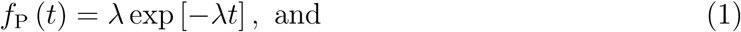

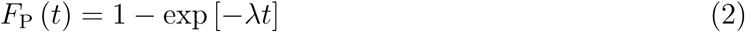

where *t* is the waiting time between events—that is, the time difference between the start of consecutive events. The average number of events in a given interval can be calculated from the rate parameter as 1*/λ*. The waiting time *t_i_* for *i* number of events from a total of *N* number of events is:^53,54^

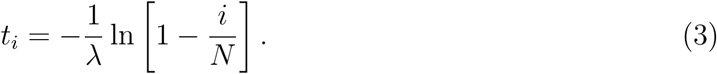

Equation 3 is used to generate a linear expression between the event waiting times *t_i_* and the distribution of events, calculated using the logarithm of ratio of remaining events ln [1 − *i/N*]. The slope of this linear relation is the rate parameter *λ*, inverse of which calculates the number of events in a given time *T*. This method was used to assess whether the distribution of ion crossing events, as defined above, and the rate of ions crossings through the channel conform to a Poisson process.

To account for the deviation between the observed distribution of waiting times between events and the expectations of a single Poisson model, we employed a chained double-Poisson process. This model incorporates two coupled Poisson process with two independent rate parameters; *λ*_lag_, depicting the lag time in events and *λ*_cond_, which defines the rate of conduction. Further details of physical foundation of this method are provided in the Results section and in the Supporting Information.

The PDF and CDF of a double-Poisson distribution are expressed as:

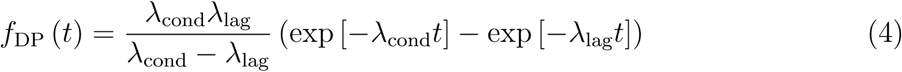

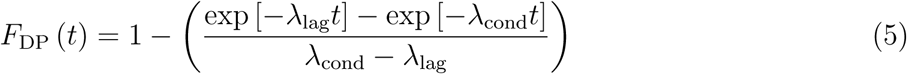

Derivation of these expressions are found in SI, section S3.

## Results and Discussion

### Single-channel recordings

To gain a meaningful basis of comparison for our molecular simulations, we recorded single-channel currents from the (cation-selective) human wild-type α7-nAChR. To this end, we expressed this Cys-loop receptor heterologously in HEK-293 cells and performed patch-clamp recordings in the cell-attached configuration. Currents were elicited by extracellular ACh (1 or 100 µM), and openings were prolonged by PNU-120596 (3 µM), a TMD-binding ligand that slows down desensitization^55^; in the absence of the latter, openings are exceedingly short, and thus, the single-channel conductance is prone to be underestimated. Under these conditions, and in the presence of ∼155-mM K^+^ in the pipette (and the nominal absence of divalent cations), the single-channel conductance was ∼193 pS for inward currents and ∼205 pS for outward currents (Fig. 2, top panel). In the presence of ∼155-mM Na^+^ in the pipette (and the nominal absence of divalent cations), instead, the inward conductance was ∼109 pS (Fig. 2, top panel). These values represent the conductance of the fully open current level (Fig. 3).

**Figure 1:**
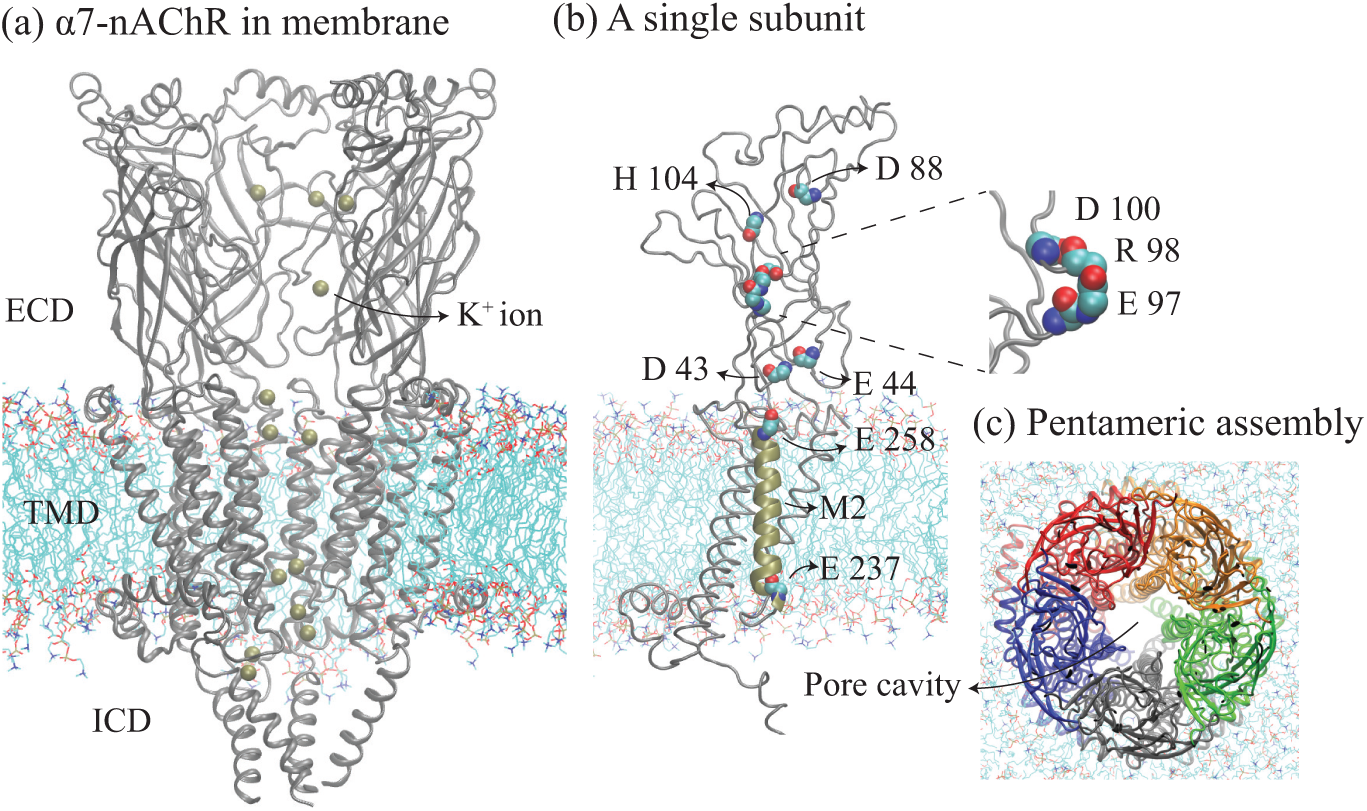
Snapshots from the simulation of PDB ID 7KOX: (a) the assembly of the α7-nAChR in the membrane bilayer. For clarity, only four of the five subunits are shown. The panel in the middle (b), shows a single α7 subunit with charged residues (and His 104) found on the lining of the pore. One of the M2 α-helices, which faces the aqueous lumen of the transmembrane pore, is thickened (colored in zinc). The loop with Asp 100, Arg 98 and Glu 97 is enlarged to the right. A top view of the pentameric assembly of the α7-nAChR is shown in panel (c) where each subunit is colored differently.

**Figure 2:**
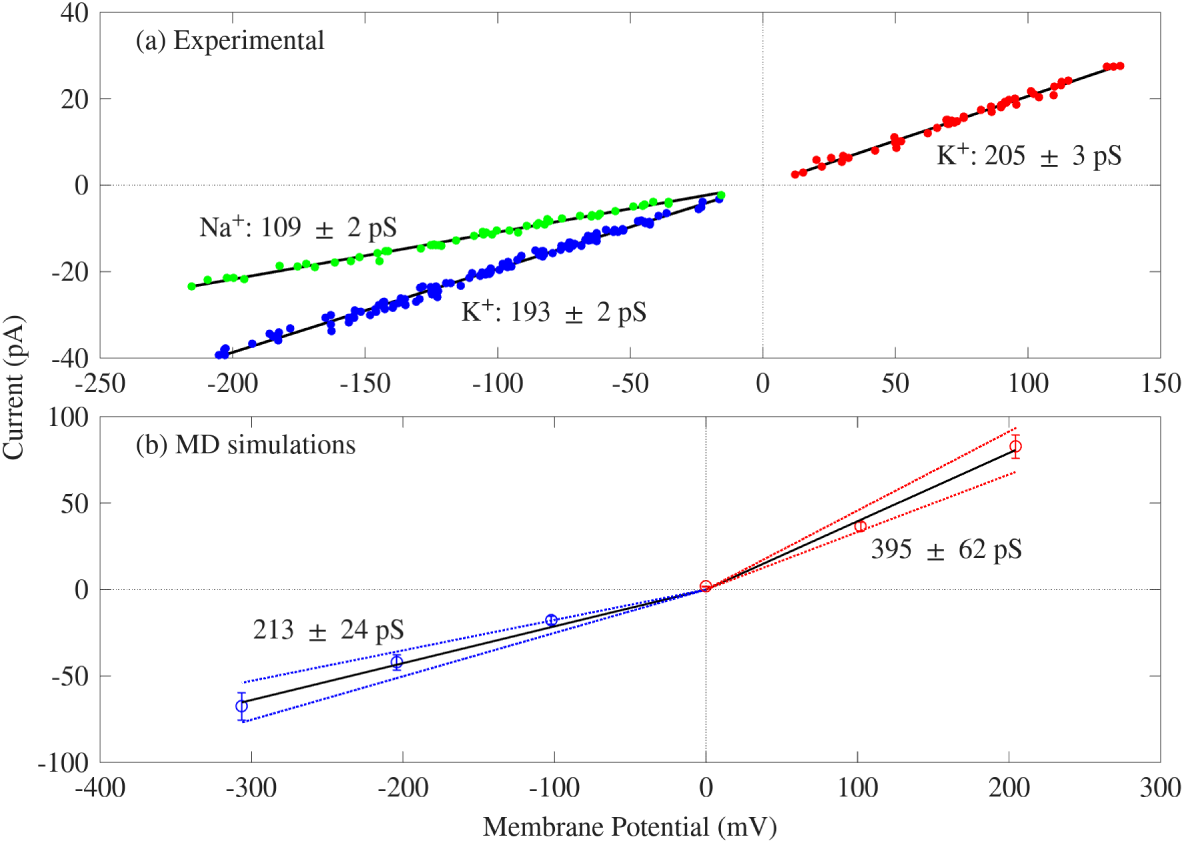
Average currents measured (a) experimentally and (b) using MD simulations at different voltages. Blue scattered points in panel (a) are K^+^ inward-current amplitudes recorded from 10 separate cell-attached patch-clamp experiments; red scattered points are K^+^ outward-current amplitudes recorded from 6 separate cell-attached patches; and green scattered points are Na^+^ inward-current amplitudes recorded from 7 separate cell-attached patches. The fully open current level was markedly less well-defined for outward than for inward currents, making the corresponding conductance estimate less certain. The (non-zero) intrinsic membrane potential of each patched cell was accounted for by adding its approximate value to the pipette potential. Marked circles in (b) are average currents calculated using (unrestrained) simulations of 7KOX in KCl solution at the given voltages (error bars represent standard errors). Two solid black lines are the fitted linear current-voltage relations at negative (blue) and positive (red) potentials, separately, with upper and lower 95% confidence (dashed lines). The slopes in each case give the single-channel conductance.

**Figure 3:**
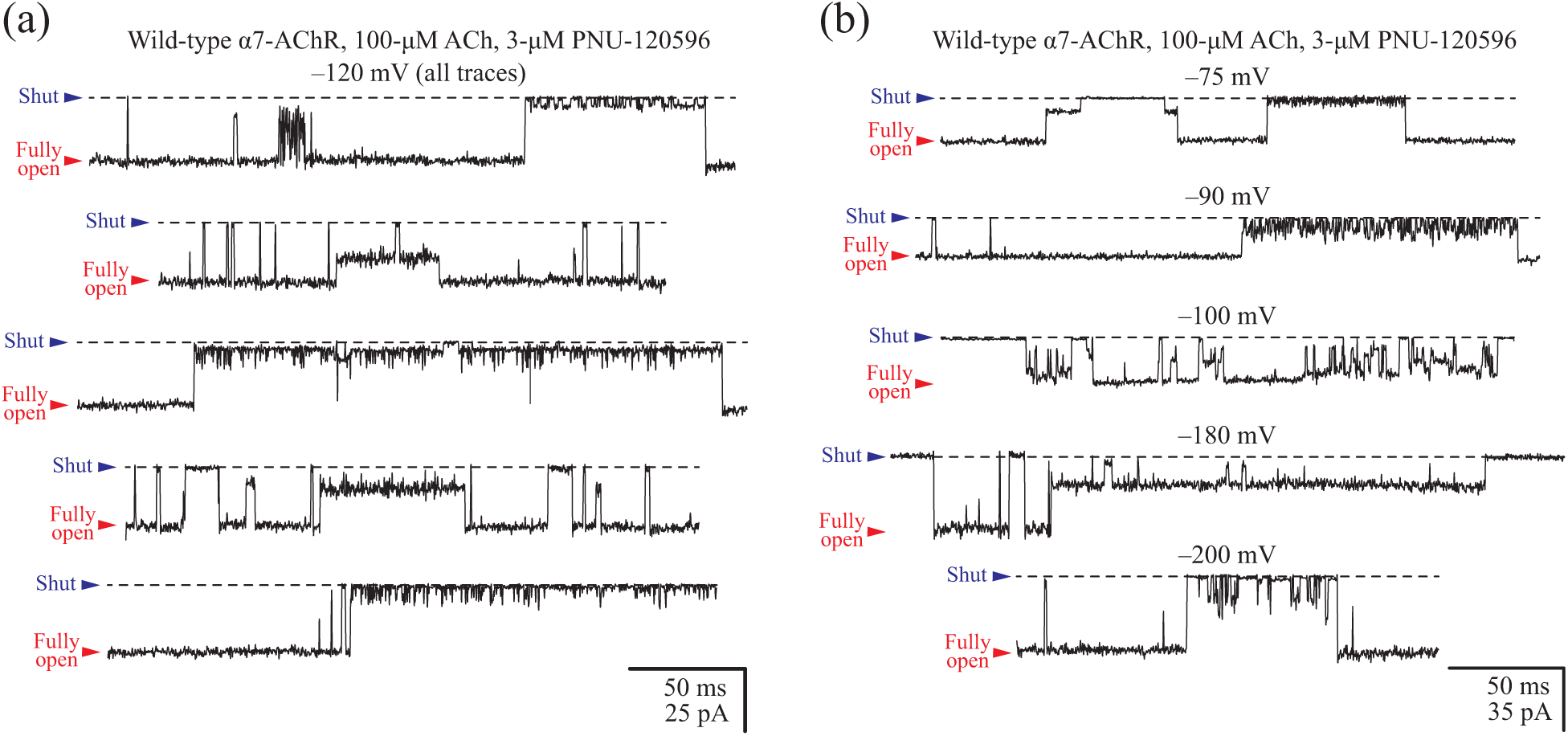
Single-channel K^+^ currents from the human α7-nAChR in the presence of a positive allosteric modulator. (a, b) Single-channel activity recorded in the cell-attached configuration from the human α7-nAChR heterologously expressed in HEK-293 cells. The positive allosteric modulator PNU-120596 was present in the ∼155-mM K^+^ pipette solution at a saturating concentration (3 µM). Under these conditions, current sojourns in partially open levels of conductance were frequent and, often, very long lived. The indicated electrical potentials correspond to those applied to the pipette; openings are downward deflections; and—for display purposes—the bandwidth was narrowed to DC–5 kHz. Black dashed lines denote the zero-current baseline. The displayed current traces were recorded from different cells.

Notably, currents through the human α7-nAChR dwelled on partially open levels (Fig. 3) with a much higher probability and for much longer durations than do currents recorded through, for example, the adult-muscle AChR^56^ (a heteropentamer of (α1)_2_β1δε subunit composition). These current sublevels were difficult to characterize quantitatively because they seemed to occur essentially anywhere between the fully open and the shut levels. In other words, these sublevels could not be assigned to discrete subconductance classes. These partial openings may reflect incomplete degrees of pore expansion, protonation–deprotonation events and/or changes in the rotameric state of side chains; the reasons why they are so prominent and their amplitude is so variable in the α7-nAChR remains puzzling. Regarding the single-channel conductance of inward currents carried by K^+^, we note that the value of ∼193 pS is remarkably close to the value of ∼185 pS measured from the mutant adult-muscle nAChR engineered to also contain five glutamates at the intracellular end of the pore^56^ (position –1’; Glu 237 in the α7-nAChR; the wild-type muscle nAChR has four glutamates and one glutamine at this position). Moreover, the ratios of K^+^-to-Na^+^ conductances of these two channels closely agree with each other (∼1.8 for the α7-nAChR and ∼1.6 for the muscle nAChR; results included in SI, section: S4). It follows that the pores of these two closely related Cys-loop receptors have very similar electrostatic properties.

### Comparison of Simulation Results on PDB structures 7KOX, 7EKT, 8V80, 8V82, and 9LH5

To simulate ion conduction through the α7-nAChR, we compared MD simulations of the different structures with PDB IDs 7KOX, 7EKT, 8V80, 8V82, and 9LH5. As described in the methods section, for each of the five atomic models, we performed three sets of different simulation runs. These MD runs are all in 150 mM concentration of KCl solution. The first set is composed of 500-ns long simulations of systems where the structures of the channel were restrained to their published PBD states. The second set consists of five unrestrained molecular dynamics runs simulated without the ligands. The third set consists of five, also unrestrained MD runs simulated with the ligands modeled in the PDB structures. The summary of conductances calculated from each set of runs is presented in Table 2. Results from each simulation are documented in Table 3, and Tables S10 to S16. The average channel radii of these structures are shown in Fig. 4 and the electrostatic profiles, densities of K^+^ ion, water densities, and hydration of ions can be found in SI, Fig. S4.

**Figure 4:**
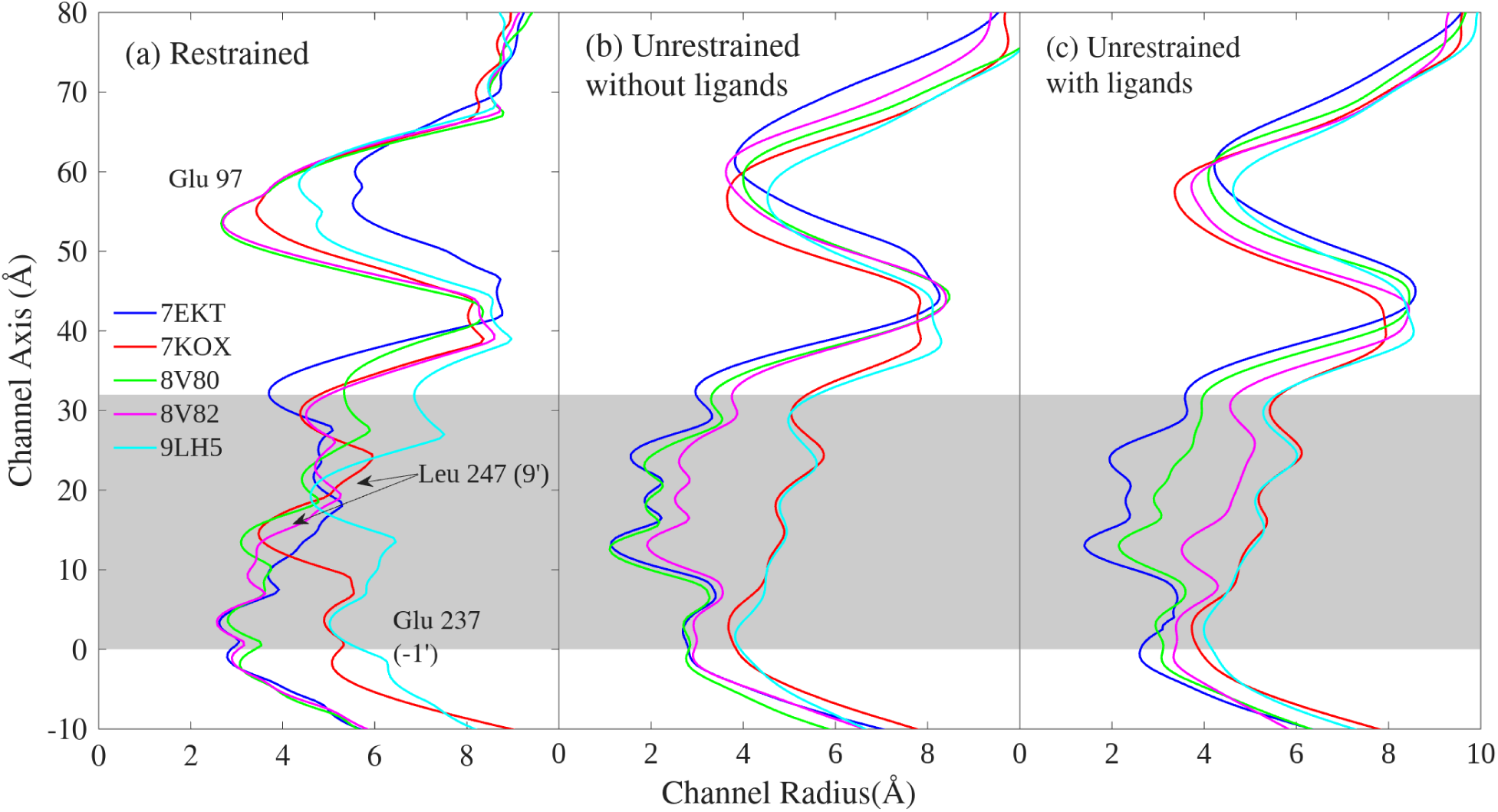
The average profiles of the five simulated structures; results are averages from all the runs. The gray area indicates the TMD. The zero value along the vertical axis corresponds to position Glu 237 (–1’).

**Table 2:** Summary of conductances (all values are in pS) calculated during restrained and unrestained simulations in 150 mM KCl solution, both with and without their respective ligands in designated structures. All results reported here are with a membrane potential of V*_e_* = −102 mV. Results are averages from all MD simulations ran for different durations (see next sections and SI, section S5 for details).

| PBD IDs | Restrained<br>Without Ligands | Unrestrained<br>Without Ligands | Restrained<br>With Ligands |
| --- | --- | --- | --- |
| 7EKT | $134 \pm 11$ | $0 \pm 0$ | $0 \pm 0$ |
| 7KOX | $12 \pm 5$ | $219 \pm 22$ | $182 \pm 24$ |
| 8V80 | $24 \pm 9$ | $0 \pm 0$ | $57 \pm 40$ |
| 8V82 | $123 \pm 10$ | $49 \pm 43$ | $225 \pm 23$ |
| 9LH5 | $128 \pm 17$ | $419 \pm 44$ | $431 \pm 21$ |

**Table 3:** Summary of events at V*_e_* = −102 mV from unrestrained simulations of 7KOX without bound to ligands. Simulations ran longer are broken in intervals of 200 ns. The first four columns from the left, after indices of the runs, are TB and BT event counts from all simulations for K^+^ and Cl*^−^* ions as marked. The net events, TB_K+_ + BT_Cl_*_−_* − BT_K+_ − TB_Cl_*_−_*, are used to calculate the conductance, in pS, which are given in the last 2 column. The average of quantities in each column is given in the last row of the table with standard error.

| Index | TB<sub>K<sup>+</sup></sub> | BT<sub>K<sup>+</sup></sub> | TB<sub>Cl<sup>-</sup></sub> | BT<sub>Cl<sup>-</sup></sub> | Net Events | Conductance [pS] |
| --- | --- | --- | --- | --- | --- | --- |
| 1 | 50 | 17 | 0 | 0 | 33 | 260 |
| 2 | 40 | 17 | 0 | 0 | 23 | 181 |
| 3 | 32 | 20 | 0 | 0 | 12 | 94 |
| 4 | 39 | 27 | 0 | 0 | 12 | 94 |
| 5 | 31 | 21 | 0 | 0 | 10 | 79 |
| 6.a | 45 | 12 | 0 | 1 | 34 | 264 |
| 6.b | 45 | 16 | 0 | 0 | 29 | 225 |
| 7.a | 32 | 15 | 0 | 1 | 18 | 142 |
| 7.b | 28 | 6 | 0 | 0 | 22 | 174 |
| 8.a | 40 | 23 | 0 | 0 | 17 | 134 |
| 8.b | 62 | 16 | 0 | 0 | 46 | 362 |
| 9.a | 51 | 18 | 0 | 0 | 33 | 260 |
| 9.b | 43 | 13 | 0 | 1 | 31 | 244 |
| 10.a | 47 | 16 | 0 | 0 | 31 | 242 |
| 10.b | 39 | 10 | 0 | 1 | 30 | 234 |
| 10.c | 62 | 13 | 1 | 1 | 49 | 383 |
| 10.d | 65 | 14 | 0 | 0 | 51 | 399 |
| 10.e | 43 | 21 | 0 | 0 | 22 | 171 |
| Average |  |  |  |  |  |  |
| - | 44 ± 2 | 16 ± 1 | 0 ± 0 | 0 ± 0 | 28 ± 3 | 219 ± 22 |

The left panel of Fig. 4 shows the results of the radii from the restrained simulations indicating the narrowest position at Leu 247 (position 9’) if 7KOX and 9LH5, whereas 7EKT, 8V80, and 8V82 have the narrowest constriction at Glu 237 (position -1’). During the restrained simulations, 7KOX frequently exhibited transient breaks in the water column at the Leu 247 constriction. 9LH5 has a wide pore throughout, allowing multiple ion crossings in both directions (see Table S10), and among the five restrained structures, it was the most hydrated (see Fig. S4). Atomic model 8V80 has two constrictions, one at Glu 237 and another one at Leu 247 (position 9’), which likely contributed to the low conductance computed during the simulation. 8V82 and 7EKT could be approximated to have a ‘V-shaped’ transmembrane pore. These two models displayed higher conductances likely because of the negatively charged glutamate (Glu 237) at the position of the constriction, allowing ions and water to pass, even if overall narrower than 7KOX and 9LH5. Although all five restrained structures allowed K^+^ permeation, the rate of ion conduction does not match the experimentally obtained value.

The unrestrained simulations revealed that quickly upon removing restraints, the con-ducting and non-conducting profiles distilled out. Interestingly, there is almost no distinction between the TMD profiles of 7KOX and 9LH5 during unrestrained runs, even though they start from two PDB structures with distinct TMD PDB profiles. In these two models, the side-chains of the lumen-facing residues, particularly Leu 247 (9’), move away from the pore, widening the channel. These two modeled structures feature a similar ‘V-shaped’ pore profile with the narrowest position at Glu 237 (-1’). A comparison of pore-radius profiles of these two models (which are almost identical at the level of the TMD) reveals that the largest difference occurs at the extracellular vestibule’s Glu 97 position; 9LH5 is much wider here, which could explain why the conductance computed for this channel is considerably higher than 7KOX’s.

We also compared simulations performed with and without the ligands modeled in the cryo-EM structures. The effects of ligand retention differed substantially across models. For 8V82, we found that the presence of ligands was essential for maintaining a conductive pore. Simulations without these ligands produced permeation in only one run (see Table S15), whereas the simulations of ligand-bound models yielded conductances consistent with experiment (see Table S11). In contrast, the 7KOX and 9LH5 models do not have PNU-120596 in the PDB structures, and these simulations showed similar permeation properties with or without their resolved orthosteric agonists, indicating that they possess an intrinsically stable conductive geometry. The behavior of 7EKT differed markedly. Although the atomic model also contained PNU, the modulator did not remain stably bound during unrestrained simulations. In multiple trajectories, at least one molecule exited its binding pocket, while others rotated substantially within it. This loss of stability of the modulator could likely explain why the pore collapses under otherwise comparable conditions. These observations are consistent with a previous MD study^16^, which reported that the modulator adopts a flipped orientation in this structure relative to its binding pose in 8V82, reducing its ability to stabilize an open conformation. The 8V80 model has TQS as the TMD-bound ligand. Although TQS remained bound throughout all simulations, only two out of five trajectories exhibited measurable conduction (see Table S12). Across all unrestrained conductive states, regardless of the presence or absence of ligands, we observed that the channel consistently relaxed into a funnel-like "V-shaped" pore profile in the TMD, with Glu 237 (-1’) forming the narrowest constriction.

To summarize, we found all the models to be in an open-state during the restrained MD simulations, but the conductance calculated (Table 2) from any of these structures did not coincide with the results obtained from the experiments. During the unrestrained MD simulations of the ligand-free atomic models, only 7KOX and 9LH5 showed conduction through the channels; 7EKT (always), and 8V80 and 8V82 (for the most part) remain non-conductive throughout the runs (SI, section S5.4). We found 7KOX, 8V82, and 9LH5 to be conductive in all the MD runs; model 8V80 was conductive in only a subset of the MD runs; whereas 7EKT was non-conductive in all of the ligand-bound unrestrained runs (SI, section S5.3). We would like to emphasize that the pore-radius profiles of the atomic models and their unrestrained MD-simulated counterparts differ significantly for all five models (Fig. 4). For example, the narrowest constriction of the TMD pore of 7KOX and 9LH5 moves from the ring of Leu 247 side chains halfway through the pore (position 9’), in the PDB models, to the intracellular mouth, around Glu 237 (position –1’), upon unrestrained simulation. We found that the conductance obtained from the unrestrained simulations of the ligand-bound 7KOX and 8V82 best matched our experimental measurements. Results from ligand-free runs of only the 7KOX model were found to be in close agreement with the experimentally obtained single-channel results. The conductance computed for ligand-free 9LH5 model was two-fold higher, likely due to a wider extracellular constriction compared to that of 7KOX. Notably, the presence of orthosteric ligand (the agonist epibatidine) did not significantly alter the conductive geometry or permeation statistics relative to the ligand-free model, indicating that MD-simulated 7KOX possesses an intrinsically stable open conformation. Since the aim of the our study was to understand the mechanism of ion conduction through the open state of the α7-nAChR, the discrepancies among the non-conductive structures were not further explored. Because both ligand-bound and unbound models relax into the same conductive state, we focused our detailed mechanistic analysis on the ligand-free 7KOX model. The latter provides a minimal system for examining the intrinsic features of ion permeation. For comparison, we also analyzed permeation statistics for 8V82 under unrestrained conditions with its resolved ligands retained, since this model also produced close agreement with experiment.

### Current–voltage characteristics

The obtained (single-channel) current–voltage (I-V) plots from the MD simulations in KCl solution are shown in Fig. 2, panel (b). The values of the membrane potential, the length of each simulation, and the number of replicas are summarized in Table S8. The slopes of the linear fits were used to calculate the single-channel conductance. One batch of five MD simulations were conducted at 22*^◦^*C in KCl solution, to match the temperature of the experiments, and results from these are presented in Table S9. Besides these, all simulations were conducted at 30*^◦^*C, as discussed in the methods section. Results from MD simulations in NaCl solution are included in Table S14.

For inward currents, the calculated single-channel conductance of K^+^ ions (213 ± 24 pS) agrees closely with the experimentally obtained value (193 ± 2 pS). The conductance calculated (175 ± 19 pS) from simulations at 22*^◦^*C is lower than those at 30*^◦^*C, yet still consistent with the results from experiments. This decrease in conductance is expected forall the models at lower temperature. For outward currents, the calculated single-channel conductance was 395 ± 62 pS, nearly twice as large as the experimental estimate (205 ± 3 pS). The discrepancy is not clear, but we surmise it may be due to the absence of the full intracellular domain in available structural models, the current-blocking effect of cytosolic Mg^2+^ in patch-clamp cell-attached recordings of outward currents, or the lower occupancy of the fully open state at positive potentials. In our simulations, the ratio of outward to inward conductance was ∼2.

For the Na^+^ inward currents, we also found a close agreement between the results from the MD calculations (128 ± 11 pS) and the experimentally measured conductance value (109 ± 2 pS). Previous computational electrophysiology estimates of the Na^+^ conductance of this receptor channel in the inward direction under the same ion conditions (symmetrical 150-mM NaCl) yielded values of 78 pS^12^ and 44 pS^11^. Although our MD runs for Na^+^ were conducted at a single voltage of ∼100 mV, these simulations captured a ratio of K^+^-to-Na^+^ conductance of ∼1.7, in remarkable agreement with our experimental observations (∼1.8; see section: single-channel recording).

### Channel Profile and Electrostatics

The positions of ionizable residues (*P_r_*) along the pore, the channel radius (*R*_0_), the electrostatic potential (Φ*_e_*), the ion densities, and the hydration number of ions along the channel pore are plotted in Fig. 5.

**Figure 5:**
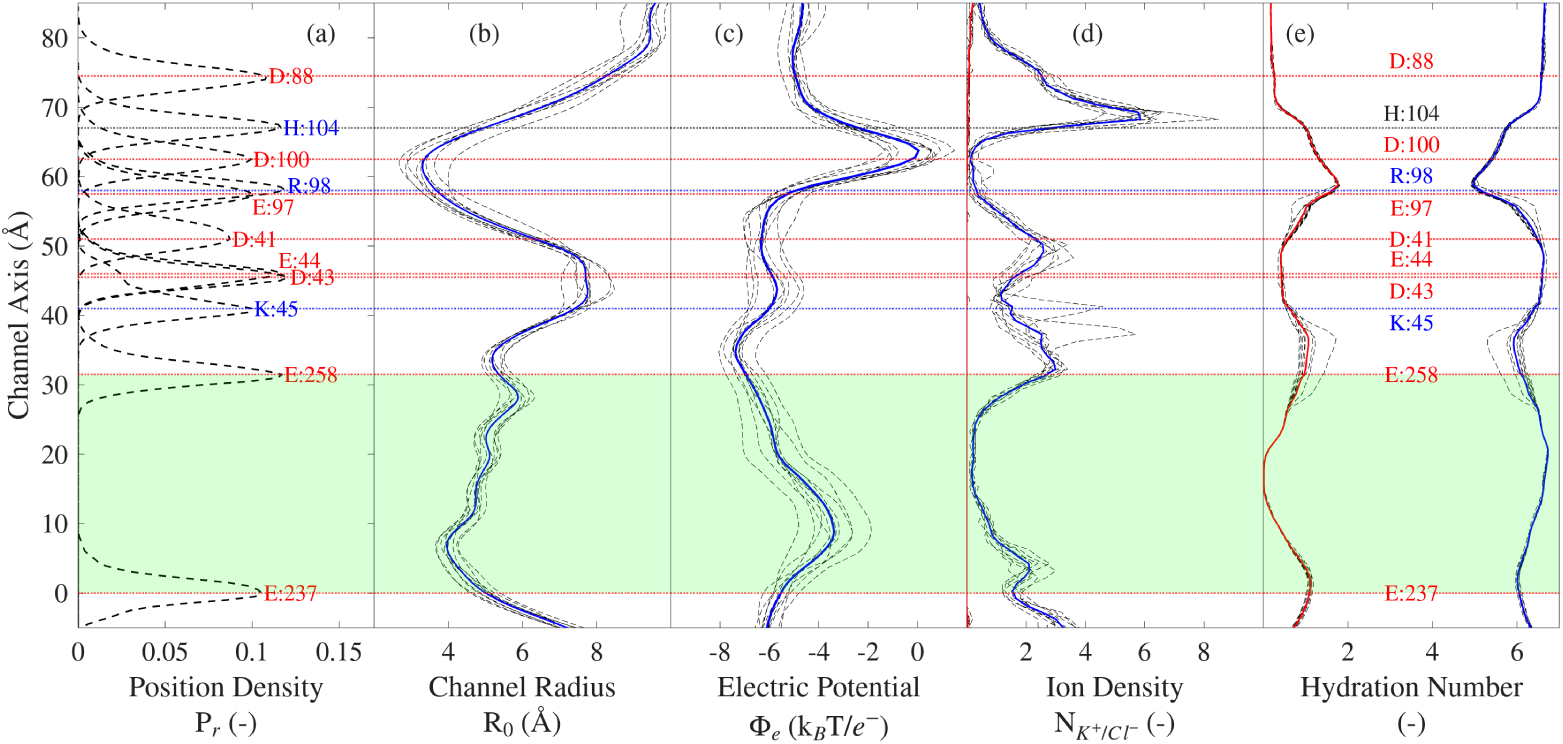
Channel profile of the α7-nAChR (7KOX) during the unrestrained MD simulations (Table 3). For reference to the positions of residues labelled here, a single α7-AChR subunit is shown in Fig. 1 (b). Along the vertical axis, zero is assigned to the average positions of Glu 237 (-1’), and the TMD region is shaded in green. Panel (a) is the position density of ionizable residues, (b) is the variation in channel radius, (c) is the electrostatic potential map of the channel, (d) shows the density of K^+^ (in blue) and Cl*^−^* ions (in red) normalized to their concentrations in bulk water. Panel (e) is the hydration number of K^+^ ions inside the channel (in blue) and the number water molecules replaced by oxygen atoms of protein residues (in red).

The position density in Fig. 5 (a) shows the normalized probability of finding ionizable residues along the inner lining of the pore (the pore axis of the channel). These were obtained by considering all atoms (including hydrogen) in all 5 subunits, calculated from simulated data sampled every 100 ps.

The radius of the channel pore was calculated using HOLE (with default van der Waals radii of atoms) every 100 ps at vertical intervals of 0.5 Å for each simulation (Fig. 5 (b)). The average radii from individual runs are plotted as dotted lines, and the solid blue line represents the radius of the channel averaged over all runs. The narrowest constriction of this particular model (unrestrained, MD-simulated 7KOX) occurs between Arg 98 and Asp 100, in the extracellular domain; the second narrowest constriction occurs at the intracellular end of the pore, at position Glu 237.

The electrostatic potential along the channel pore, Φ*_e_*, was calculated incorporating the smooth partial-mesh Ewald method (PME) using the PMEPot plugin in VMD. These calculations take into account all charges in the simulation box, which include ions and charged residues of the protein. The ion density was obtained by counting the number of ions in 0.5-Å-thick disks of radius 17.3 Å along the vertical axis of the pore. The plotted density was normalized to the density of ions in 150-mM KCl in water, which was ∼0.1 ions in this volume of the disk. The potential and number of ions were measured from frames saved every 100 ps, and the average of each run was plotted as dashed lines in Fig. 5 (c) and (d) respectively; the solid blue line represents the average over all 10 runs in each case. The individual result of each run deviates from the average, which reflects the variation in charge distribution between different runs. The highest potential barrier faced by ions moving along the longitudinal axis of the pore occurs near the narrowest constriction, in the ECD.

Fig. 5 (e) shows the hydration number, that is, the number of water molecules in the first hydration shell of K^+^ inside the channel. The threshold radius for this calculation (3.52 Å) was given by the radial density function of K^+^ created using a water-box MD simulation with the same ion concentration (150-mM) as that used in the simulations of ion permeation.

The solid blue curve is the average number of water molecules in the first hydration shell, and the dotted lines show values from each simulation run. The solid red curve is the average number of water oxygen atoms replaced by oxygens from protein residues; the dotted lines show values from each run. Oxygen atoms of protein replace those of water most extensively at the narrowest constriction of the pore (residues Glu 97 and Arg 98, in the ECD), followed by the rings of glutamates at either end of the transmembrane pore (Glu 237 and Glu 258). At most, K^+^ loses two water molecules from the first hydration shell (out of an average of six) as it permeates during the MD simulation.

### Event Distribution and Statistics

The distribution of events from all replicas is summarized in Table 3. The indices of the Table represent individual MD runs. Any simulation longer than 200 ns was broken down into sections of 200-ns, and these are further identified by Roman letters. For example, run 6 was 400 ns long, and 6.a and 6.b are considered independent runs of 200 ns. Event counts — that is, the first four columns of the Table after indices — are either positive (TB_ion_), when an ion enters from the ECD (Glu 237) and exits into the ICD (Glu 258), or negative (BT_ion_), when an ion crosses the TMD in the opposite direction. The second-to-last column of this table gives the net number of events, defined here as the net translocation of charges through the transmembrane pore in either direction. The table reveals a significant variation in the number of net events, and hence conductance, between different MD runs, with the overall conductance ranging between 79 and 399 pS. This indicates that multiple measurements are needed to correctly estimate the mean conductance, as individual measurements could result in a large error. To estimate the variability of the individual measurements relative to the mean, we calculate the ratio of standard deviation to average number of TB_K+_, and BT_K+_ events to be 0.24 and 0.33 respectively, suggesting that the ion conduction in the TB direction is more consistent. The distribution of event durations is shown in Fig. S10 for ion translocations in either direction. These probability densities were fitted with a log-normal distribution (dashed lines on the figure), the mean duration of events being ∼1.24 ns for both inward and outward events. We calculated the diffusion coefficient of K^+^ in the TMD pore of the channel to be 0.36 m^2^/s, that is, ∼0.2 times the rate of free diffusion of this ion in water along a single dimension^57^. The average event duration calculated for ions crossing the TMD of 8V82 bound to ligands is slower than this, see Fig. S10, where an average time for a TB*_K_*_+_ and BT*_K_*_+_ event is ∼2.37 ns and ∼1.84 ns, respectively.

To ascertain whether the differences in the number of events between different runs stem from conformational differences of the channel, we performed several types of analyses:

1. Computing the average structures and calculating the RMSD between these structures (results in Fig. S7);
2. Performing principal component analysis, PCA (results in Fig. S8);
3. Performing *t-*distributed stochastic neighbour embedding, tSNE (results in Fig. S9).

The results show that these analyses failed to identify a correlation between computed single-channel conductances and structural features. A brief overview of these analyses are included in the SI, section S9.

Next, we modelled the distribution of ion-crossing events with a temporal Poisson process. Fig. 6 shows the probability density function of waiting times (PDF*_w_*) between consecutive events. Waiting time is the time interval between the start of two consecutive events. This analysis was performed for K^+^-ion crossing events only, and the distributions of crossings in either direction are shown separately.

**Figure 6:**
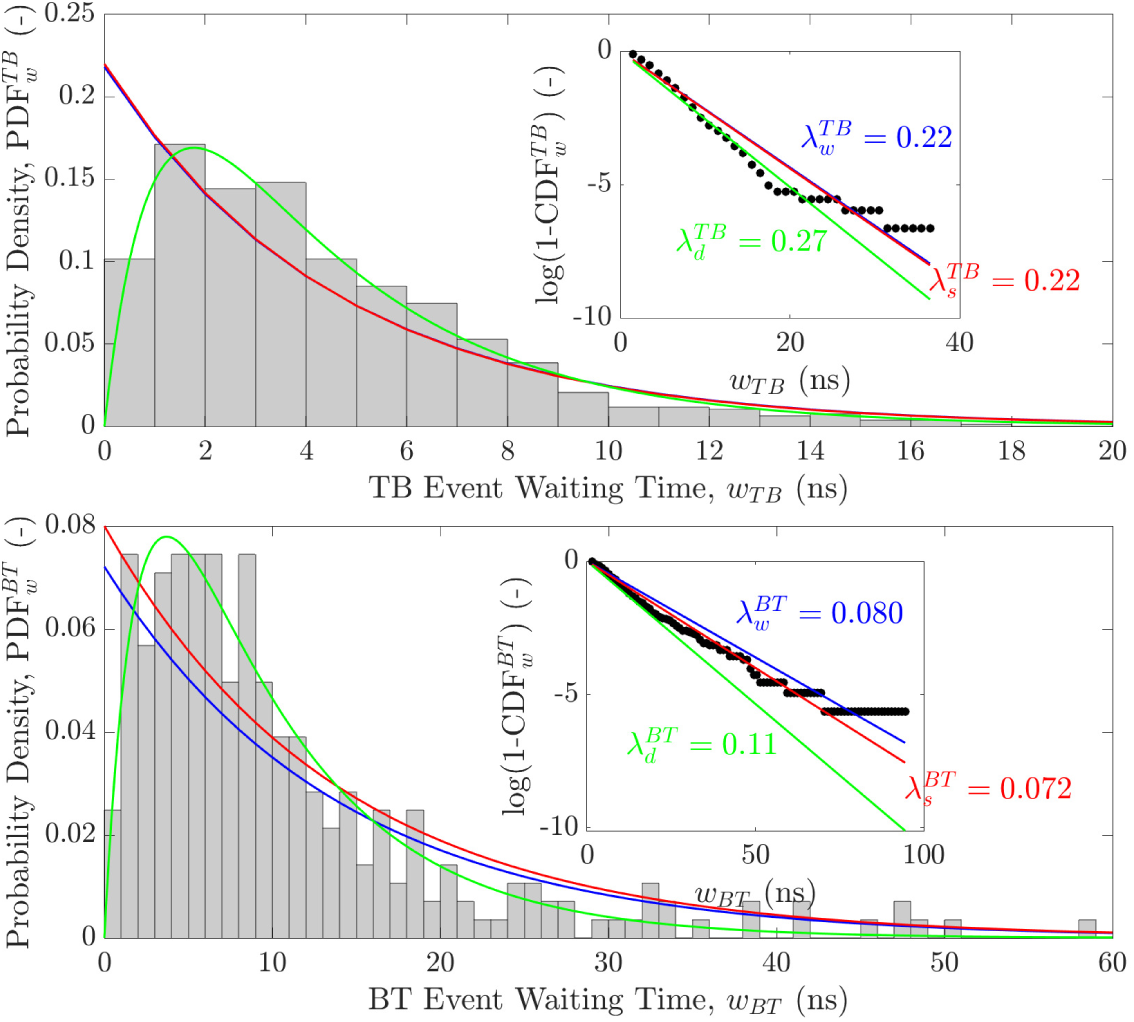
Probability density distributions of waiting times between consecutive TB*_K_*_+_ (upper panel) and BT*_K_*_+_ (lower panel) events. Data fitted to a pure Poisson (in blue, eq. 1) and double-Poisson process (in green, eq. 4) from all 10 simulations (see Table 3) at V*_e_* = −102 mV. The linear fits to the data using eq. 3 are show in inserts for each plot. The red curves are plots with the rate coefficients estimated from average number of events calculated from the simulations (see Table 3). The blue curves are calculations from the linear fit. The green curves are double-Poisson process, the slope on the linear fit does not account for the lag time. The rate coefficients are as shown on the plots, for TB events these are the same (the red plot is overlapping the blue plot).

The blue line on the plots shows an exponential decay with Poisson parameters 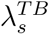 = 0.22 ns*^−^*^1^ and 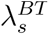 = 0.072 ns*^−^*^1^ obtained by fitting the data to a Poisson distribution. The linear regressions were forced to intercept the y-axis at 0, which is the shortest possible waiting time. The solid red lines are the exponential decay with Poisson parameter calculated from the average number of events 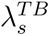 = 44*/*200 = 0.22 and 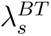 = 16*/*200 = 0.08 (events/ns). The inserts of the figure show the plots of the distribution based on the expression in eq. 3; the blue and red lines are linear fits with slopes 1*/λ_w_*and 1*/λ_s_*respectively, both passing through the origin.

To better reconcile the discrepancy between the predictions of a pure Poisson process—which allows arbitrarily short inter-event intervals—and the observed delay between consecutive conduction events in simulations, we examined individual ion trajectories. Visual inspection suggests that K^+^ ions are temporarily delayed when traversing regions near the charged residues at the termini of the transmembrane domain (TMD). These charged residues are surrounded by additional K^+^ ions which appear to electrostatically repel incoming cations. For example, an ion initiating a top-to-bottom (TB) event near Glu-258 may transiently occupy a position that inhibits subsequent ions from entering the pore. As a result, the next ion must wait until the preceding one has moved sufficiently deeper into the TMD, introducing a delay not captured by a simple Poisson process. To model this behaviour, we employed a double-Poisson process, comprising two sequential Poisson-distributed waiting times: the first accounts for the initial lag due to inter-ion electrostatic interference, and the second models the stochastic interval between conduction events themselves.

To model the distribution of waiting times using the double-Poisson process, we optimized the rate parameters, *λ*_lag_ and *λ*_cond_, in eq. 4, governing this model to fit the simulation results. The resulting optimized distributions are plotted in green in Fig. 6. The lag times in this model for TB and BT events are 1 ns and 1.53 ns, respectively, while the average waiting times are 3.62 ns and 8.14 ns.

According to a pure Poisson model, the average number of TB*_K_*_+_ events is estimated to be 44, which is identical to the average number of TB*_K_*_+_ events calculated from the 200-ns long simulation. For BT*_K_*_+_ events, the average number of events estimated from the Poisson distribution is ∼14.4 in 200 ns, and the average number of events calculated from simulations is 16. In contrast, the average number of events calculated from the double-Poisson model over 200 ns is 200*/* (1 + 3.62) ≃ 43.3 for TB*_K_*_+_ and 200*/* (1.53 + 8.14) ≃ 20.7 for BT*_K_*_+_. Overall, the agreement between these estimates is excellent.

We also modeled the event waiting time distribution from the simulations of unrestrained ligand bound 8V82 model. The event waiting time distributions are generated from events summarized in Table S11, and the results of double-Poisson model fitted to the distributions is shown in Fig. S11. The lag times here are 1.11 ns and 0.53 ns and the average waiting times are 3.32 ns and 8.14 ns for TB and BT events, respectively. The lag and waiting times for both TB and BT events between the two event distributions of 7KOX and 8V82 closely match. The noticeable discrepancy between the distribution of BT events for 8V82, as was the case for 7KOX, is mainly due to a lack of statistics (i.e., fewer number of BT events).

### Lateral Fenestrations

Lateral fenestrations have been proposed as an alternative pathway for ions to enter or exit the channel, distinct from the axial (“apical”) pathway^58^. To determine the extent to which this pathway is followed by ions permeating through the α7-nAChR, the trajectories of ions were tracked as they moved from the bulk solution into (or out of) the channel. Ions moving laterally through ICD cavities (the cytosolic “portals”) were not included in this analysis because their structure remains unsolved.

Fig. 7 shows the analysis of two TB_K+_ permeation events in blue and one BT_Cl_*_−_* permeation event in red. These are partial tracks of the ions passing through the TMD taken from one of the MD simulations (indexed 6.a of Table 3). The position of ions on the plots are marked every 100 ps, and dashed lines join them in the order in which they occurred. The middle plot shows a K^+^ permeation event that started with an excursion of the ion into the channel through the “walls” of the ECD. Visual inspection of the trajectory shows that the ion crosses the channel walls in-between two subunits, passing through the opening between the β8-β9 and β1-β2 loops. K^+^ ions taking this pathway always amount to a positive event; in other words, we did not observe any K^+^ exit the channel through these lateral openings in the ECD. Also, we did not observe any chloride ion take this pathway in either direction, possibly due to lack of chloride events.

**Figure 7:**
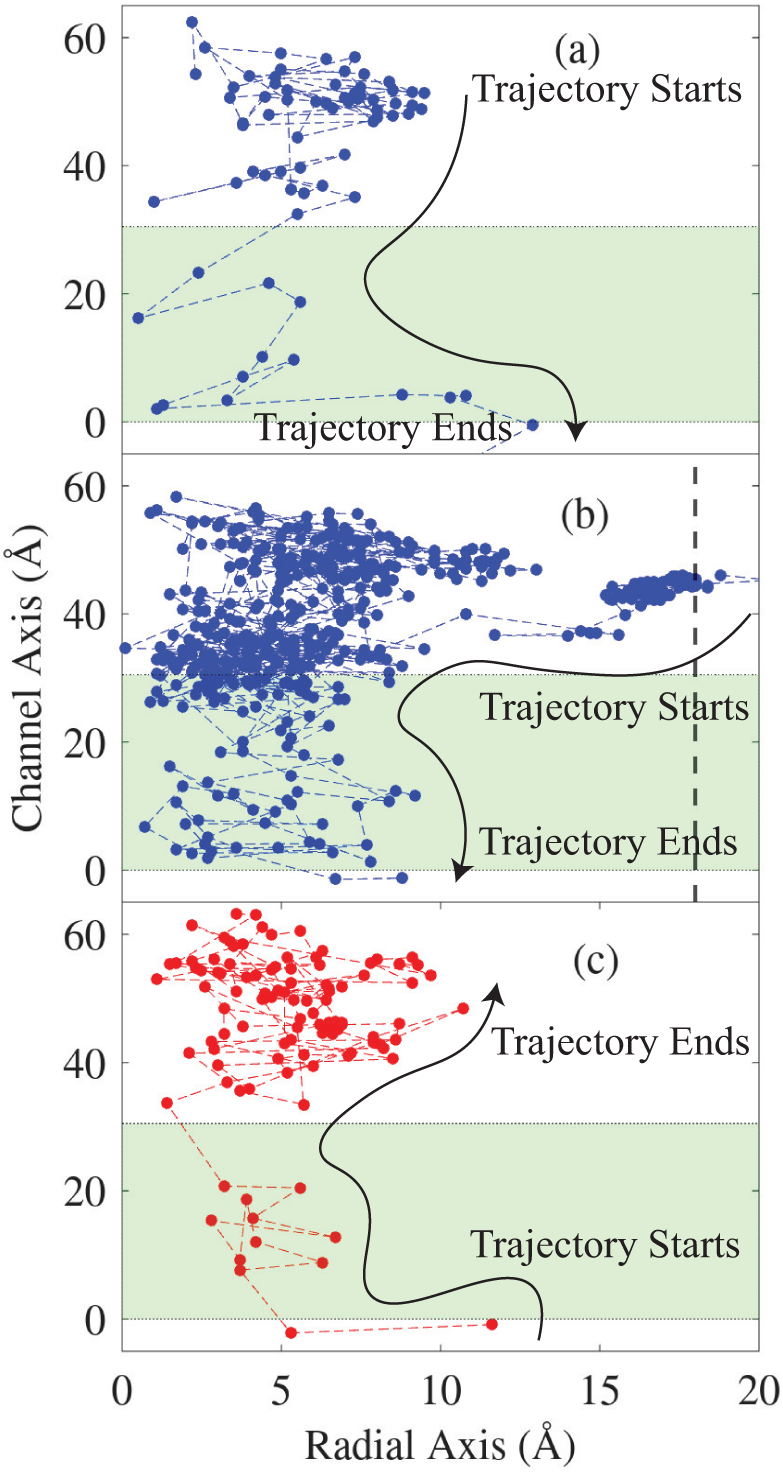
Ion movement: TB event (a), where K^+^ enters the channel axis and passes through TM; (b) K^+^ entering laterally into the ECD and moves through TMD into ICD. The vertical dashed line shows the threshold to define boundary of a laterally event. (c) Cl*^−^* entering ICD crossing TMD and out into the ECD axially. Zero on the horizontal axis is the position of Glu 237 (-1’) on the vertical axis and the TMD is shaded region.

The rate of ion permeation through these lateral fenestrations is ∼1.4 events in 1 µs. In the 10 MD runs, worth 3.6 µs at V*_e_* = −102 mV, there were a total of 5 permeation events in which the ion entered the channel through these lateral entryways. Thus, this pathway was followed by 0.6% of TB*_K_*_+_ events (of 794 total TB*_K_*_+_ events) and 0.4% of total events (of 1,089 total events in either direction). In marked contrast, in the anion-selective Cys-loop receptor α1-GlyR, MD simulations showed a much higher contribution of lateral pathways amounting to 96% of total events (717 lateral events of total 748 events in both directions) for Cl*^−^*.^58^ Despite its low probability and longer time to cross the lateral boundary, there exist clear openings between the subunits of the receptor that allow ions to enter the channel.

## Conclusions

We performed extensive MD simulations on five cryo-EM models of the α7-nAChR proposed to represent open or partially open conformations (PDB IDs 7KOX, 7EKT, 8V80, 8V82, and 9LH5). Across all models, we compared simulations conducted with the protein held restrained to the experimentally determined coordinates to simulations performed on unrestrained models whose structures were allowed to relax. All restrained simulations exhibited at least modest conductance (Table 2), but none of them yielded single-channel conductances consistent with the experimentally estimated value for the fully open state. Once the protein was allowed to relax, each model diverged from its deposited cryo-EM geometry, undergoing subtle but functionally important rearrangements of the pore-lining helices (Fig. 4 and Fig S6). These changes substantially altered ion-permeation behavior (table 2): some models lost conductance entirely, others shifted to conductances incompatible with experiment, and only two, 7KOX and 8V82, settled into stable conductive ensembles consistent with electrophysiological data (Fig. 2). These observations highlight the risk of judging the conductive/non-conductive state of an ion channel solely on the basis of their static coordinates; even modest rearrangements upon relaxation can have a profound impact on the rate of computed ion flow.

We studied the influence of bound ligands on the conductive behavior across the different α7-nAChR structures. For 7KOX, the deposited coordinates contain only epibatidine, although the experimental preparation also included the TMD-bound modulator PNU-120596. Simulations with and without epibatidine produced similar conductances, and the absence of resolved PNU did not compromise its open-state conductance. Interestingly, 9LH5, which contains only L-nicotine and was solved in the absence of any TMD-binding modulator, also relaxed into a conductive ensemble with a TM pore profile nearly identical to that obtained for 7KOX. This convergence suggests that the stable conductive geometry observed for these two structures does not require a bound modulator, even though one was present experimentally for 7KOX.

In contrast, 8V82 was solved with both epibatidine and PNU-120596, and its conductive properties depended strongly on retaining these ligands. Removing both ligands resulted in low or absent conduction, whereas including the resolved epibatidine and PNU yielded conductances in line with experiment. Because we did not simulate 8V82 with epibatidine alone, we cannot isolate the individual contributions of the two ligands; however, the similarity of 7KOX simulations with and without epibatidine suggests that PNU-120596 is likely the dominant stabilizing factor for pore conductivity in 8V82. A more modest form of ligand dependence was observed for 8V80, which was non-conductive when simulated without its resolved ligands (epibatidine and the modulator TQS), but exhibited intermittent conduction in a subset of ligand-bound simulations. Finally, for 7EKT, which contains both EVP-6124 and PNU-120596, PNU molecules migrated out of their binding pockets in a majority of simulations, and the channel collapsed into a non-conductive state. This behavior further underscores the importance of modulators in maintaining a conductive pore, and suggests that the 7EKT cryo-EM conformation does not encode a physiologically relevant stable conductive ensemble.

Among all structures tested, only 7KOX and ligand-bound 8V82 produced ensemble-averaged conductances close to the experimental single-channel value once allowed to relax (∼193 pS for inward currents in external ∼150-mM K^+^). The 9LH5 structure yielded consistently overestimated conductances, approximately twice the experimental value, even after relaxation. Strikingly, the unconstrained conductive models, i.e., 7KOX, modulator-bound 8V82, and 9LH5, converged independently onto a similar V-shaped or funnel-like pore profile, with the minimal radius positioned at the Glu 237 (-1’) site. At this location, the conserved Glu 237 residue provides a negatively charged constriction that stabilizes permeating cations. This convergence across independent starting structures suggests that the -1’-centered funnel geometry represents a structural feature of the α7-nAChR’s fully open transmembrane pore.

Inspection of the channel radii (Fig. 4, panel b and c) revealed a similar TM pore profiles of 7KOX and 9LH5, yet 9LH5 showed a markedly wider extracellular-region opening near Glu 97. This localized expansion coincided with a roughly twofold increase in conductance relative to that of 7KOX, in agreement with the known role of this region of the extracellular vestibule in the control of conductance in nicotinic receptors^59^.

For all structures, we carried out multiple independent simulations. Conductance estimates with K^+^-ion, obtained from 200 ns segments spanned a broad range (e.g., 79–399 pS for 7KOX), underscoring the need for ensemble averaging when evaluating permeation in highly dynamic open-channel conformations. To investigate possible structural determinants of conductance variability, for 7KOX simulations we analyzed a broad set of geometric and physicochemical features, including the positions of charged side chains along the pore axis, pore-radius profiles, electrostatic potentials, ion-density distributions, and hydration patterns of permeating K^+^ ions (Fig. 5). We further quantified structural variance via RMSD analysis (Fig. S7), principal component analysis (Fig. S8), and t-SNE projections across subunit copies (Fig. S9), but no single descriptor showed a consistent correlation with conductance across different runs. The permeation statistics across models remained predominantly Poissonian, with a characteristic entry lag attributable to electrostatic repulsion during ion entry. This behavior was accurately captured using a double-Poisson model, comprising linked distributions for the lag times and subsequent inter-event waiting intervals.

Interestingly, the conductive model (MD-relaxed 7KOX) also allowed ions to enter laterally through extracellular-domain (ECD) fenestrations. However, only a small fraction (0.4%) of the total ion crossings occurred via this slower lateral pathway. Because diffusion of Na^+^ is slower than K^+^, and Na^+^ ions are more hydrated, we expect that Na^+^ ions would use this entry pathway even less frequently than 0.4%, suggesting that during physiologically relevant Na^+^ influx, the majority of ions would be conducted through the central pore.

Lastly, the heterogeneous conductive behavior of these cryo-EM structures raises the possibility that they represent distinct open conformational states of the α7-nAChR. The receptor displays multiple subconductance levels in electrophysiological recordings, and the structural diversity observed across cryo-EM models may reflect a continuum of such states. Future work incorporating complete intracellular domains, resolved ligands, and additional structural intermediates will be required to fully elucidate the structural basis of α7-nAChR subconductance behavior.

## Supporting Information Available

- Supporting Information. MD simulation protocols, setup details, and ensemble parameters; methodology for channel conductance calculations; results of single-channel conductance from the muscle nAChR; summary tables of event counts and conductance results for restrained and unrestrained systems; probability density distributions of event durations and waiting times; position density and radial variation profiles of pore-lining residues; channel profiles for electric potential, ion density, and hydration; structural correlation analyses including RMSD matrices, PCA, and tSNE; and derivation of the Double-Poisson process model.

## Data Availability

- The MD simulations for study are openly available in MDposit at this link.
- Analysis scripts maybe requested by the corresponding authors.

## Acknowledgement

The authors would like to thank Dr. Richard Pastor (NHLBI, NIH) and Dr. Samuel Foley (Biophysics, JHU) for their valuable discussions. Computer simulations for this work were ran on LoBoS at (NHLBI) NIH and Rockfish at Johns Hopkins supercomputing clusters. A.D. acknowledges NHLBI grant 75N92020P00042.

This research was supported by the Intramural Research Program of the National Institutes of Health (NIH), NHLBI (HL001050) to B.B. The contributions of the NIH author(s) were made as part of their official duties as NIH federal employees, are in compliance with agency policy requirements, and are considered Works of the United States Government. However, the findings and conclusions presented in this paper are those of the author(s) and do not necessarily reflect the views of the NIH or the U.S. Department of Health and Human Services.

This research was also supported by the School of Molecular and Cellular Biology, University of Illinois Urbana-Champaign (C.G.).

## TOC Graphic

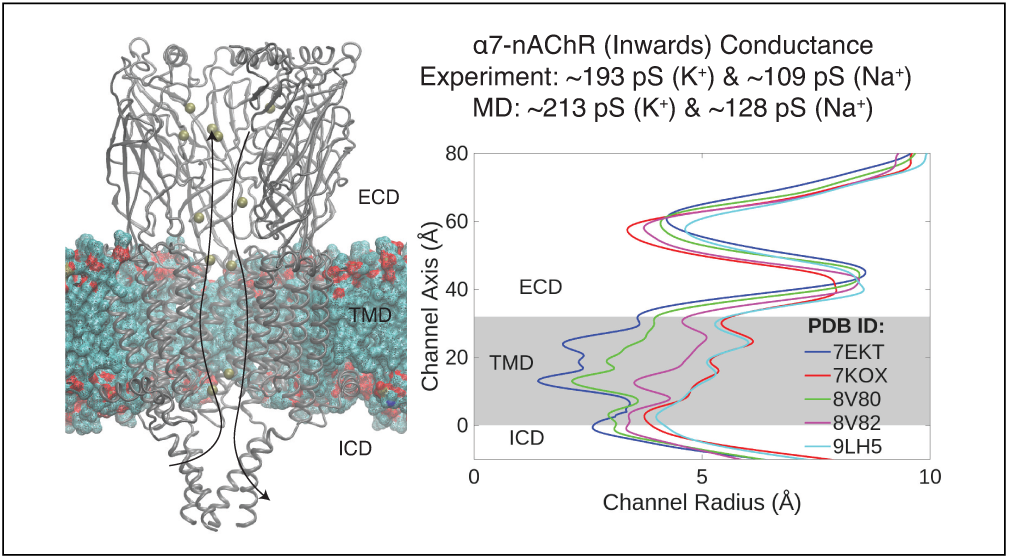

