## Supporting Information for "Characterizing the Ion-Conductive State of the *α*7-Nicotinic Acetylcholine Receptor via Single-Channel Measurements and Molecular Dynamics Simulations"

### Contents


**S1 Simulation Protocols** . . . . . **S3**
S1.1 MD Simulation: Setup Details . . . . . S3
S1.2 MD Simulation: Ensemble Details . . . . . S4

**S2 Channel Conductance: Method** . . . . . **S6**

**S3 Model of Double-Poisson Process** . . . . . **S7**

**S4 Single-Channel Conductance: Muscle nAChR** . . . . . **S9**

**S5 Channel Conductance: MD Results** . . . . . **S10**
S5.1 Simulations at 22°C . . . . . S10
S5.2 Restrained Simulations . . . . . S11
S5.3 Unrestrained Simulations with Ligands . . . . . S12
S5.4 Unrestrained Simulations without Ligands . . . . . S14

**S6 Additional Results from MD Simulations** . . . . . **S15**

**S7 Position Density of Residues** . . . . . **S16**

**S8 Variation in Positions of M2 Helix Residues** . . . . . **S17**

**S9 Correlation Analysis between Conductance and Structural Variation** . . . . . **S18**
S9.1 RMSD Variation between Average Structures . . . . . S18
S9.2 PCA and tSNE Analysis . . . . . S19

**S10 Event Duration** . . . . . **S21**

**S11 Waiting Time for 8V82** . . . . . **S22**

### S1 Simulation Protocols

#### S1.1 MD Simulation: Setup Details

| PDB IDs | Patch One Res IDs | Patch Two Res IDs | Disulfide bonds Cysteine IDs |
|---|---|---|---|
| 7KOX | Glu 1 to Leu 320 | Ile 413 to Phe 478 | 141-127 & 189-190 |
| 7EKT | Gly 23 to Lys 347 | Arg 432 to Ala 502 | 150-164 & 212-213 |
| 8V80 | Glu 1 to Lys 323 | Pro 408 to Phe 478 | 141-127 & 189-190 |
| 8V82 | Glu 1 to Lys 323 | Pro 408 to Phe 478 | 141-127 & 189-190 |
| 9LH5 | Gly 23 to Leu 343 | Pro 431 to Ala 502 | 150-164 & 212-213 |

Table S1: Summary of PDBs models in MD simulations, Res IDs numbered identical to the PDB models.

| Ensemble | Duration (ns) | GLU:44/$Ca^{+2}$ Restraint (kcal/mol/Å$^2$) | Position Restraint (Protein) (kcal/mol/Å$^2$) | Position Restraint (Lipid) (kcal/mol/Å$^2$) | Dihedral Restraint (Lipid) (kcal/mol/rad$^2$) |
|---|---|---|---|---|---|
| NVT | 0.025 | 50 | 10 | 2.5 | 250 |
| NVT | 0.025 | 50 | 5 | 2.5 | 100 |
| NPT | 0.05 | 50 | 2.5 | 1 | 50 |
| NPT | 0.1 | 50 | 1 | 0.5 | 50 |
| NPT | 0.1 | 50 | 0.5 | 0.1 | 25 |
| NPT | 2 | 50 | none | none | none |

Table S2: Summary of CHARMM-GUI 6 step equilibration protocol used before production run of unrestrained simulations. For the restrained simulations, the protein restraints are 10 kcal/mol/Å(first row, forth column). The voltage difference is 0 mV (no applied EF) during these steps. The 3rd column is applicable for PDB IDs 7KOX, 8V80, and 8V82.

| PDB IDs | Ionic Solution | Box Size (x, y, z) (Å) | $EF_z$ (kcal/mol/Å/$e^-$) | POPC Count Upper/Lower |
|---|---|---|---|---|
| 7KOX | KCl | 124×124×172 | -0.014 | 120/116 |
| 7KOX | NaCl | 102×102×185 | -0.012 | 108/100 |
| 7EKT | KCl | 124×124×172 | -0.014 | 120/125 |
| 8V80 | KCl | 128×128×192 | -0.012 | 118/131 |
| 8V82 | KCl | 98×98×194 | -0.012 | 100/105 |
| 9LH5 | KCl | 102×102×187 | -0.012 | 100/105 |

Table S3: Summary of MD simulation setup details.

#### S1.2 MD Simulation: Ensemble Details

| PDB IDs | Runs | Program | Equilibration(Duration (ns)) | Production(Duration (ns)) |
|---|---|---|---|---|
| 7KOX | 5 | OpenMM | NPT (10) | NVT (200) |
| 7KOX | 5 | OpenMM | NPT (10) | NPT (200) |

Table S4: Summary of program used for each of the unrestrained run (without ligand) in NaCl solution and simulation times at voltage $V_e = -102$ mV.

| PDB IDs | Runs | Program | Equilibration(Duration (ns)) | Production(Duration (ns)) |
|---|---|---|---|---|
| 7EKT | 3 | OpenMM | NPT (10) | NVT (200) |
| 7KOX | 1 | AMBER | NPT (10) | NVT (1000) |
| 7KOX | 4 | OpenMM | NPT (5) | NPT (400) |
| 7KOX | 5 | AMBER | NPT (10) | NVT (200) |
| 8V80 | 3 | AMBER | NPT (10) | NVT (200) |
| 8V82 | 5 | AMBER | NPT (10) | NVT (200) |
| 9LH5 | 5 | AMBER | NPT (10) | NVT (200) |

Table S5: Summary of program used for each of the unrestrained run (without ligand) and simulation times at voltage $V_e = -102$ mV.

| PDB IDs | Runs | Program | Equilibration(Duration (ns)) | Production(Duration (ns)) |
|---|---|---|---|---|
| 7EKT | 5 | AMBER | NPT (10) | NVT (200) |
| 7KOX | 5 | OpenMM | NPT (10) | NVT (200) |
| 8V80 | 5 | OpenMM | NPT (10) | NVT (200) |
| 8V82 | 5 | OpenMM | NPT (10) | NVT (200) |
| 8V82 | 5 | OpenMM | NPT (10) | NVT (400) |
| 9LH5 | 5 | AMBER | NPT (10) | NVT (200) |

Table S6: Summary of program used for each of the unrestrained run (with ligand) and simulation times at voltage $V_e = -102$ mV.

| PDB IDs | Runs | Program | Equilibration(Duration (ns)) | Production(Duration (ns)) |
|---|---|---|---|---|
| 7EKT | 1 | AMBER | NPT (10) | NVT (500) |
| 7KOX | 1 | AMBER | NPT (10) | NVT (500) |
| 8V80 | 1 | AMBER | NPT (10) | NVT (500) |
| 8V82 | 1 | AMBER | NPT (10) | NVT (500) |
| 9LH5 | 1 | AMBER | NPT (10) | NVT (500) |

Table S7: Summary of program used for each of the restrained run and simulation times at voltage $V_e = -102$ mV.

| Copies | Program | Ensemble | Replica Duration (ns) | Voltage (mV) | Current (pA) | Conductance (pS) |
|---|---|---|---|---|---|---|
| 5 | AMBER | NVT | 200 | −306 | $-67.6 \pm 7.9$ | $220 \pm 26$ |
| 5 | AMBER | NVT | 200 | −204 | $-42.1 \pm 4.5$ | $206 \pm 22$ |
| 10 | See table S5 for details | | | −102 | $-1.8 \pm 2.3$ | $175 \pm 23$ |
| 5 | AMBER | NVT | 200 | ±0 | $1.8 \pm 0.3$ | n/a |
| 5 | AMBER | NVT | 200 | +102 | $36.5 \pm 2.5$ | $358 \pm 24$ |
| 5 | AMBER | NVT | 200 | +204 | $82.7 \pm 6.6$ | $405 \pm 33$ |

Table S8: Summary of production runs (7KOX, unrestrained without ligands) at different voltages. Negative and positive voltages create inward and outward currents, respectively. The current calculated for $V_e = -102$ mV is based on all 10 runs, others are based on 5 runs.

#### S2 Channel Conductance: Method

The current through the channel was calculated as:

$$I_r \simeq \frac{1}{\Delta t} \sum_{i=1}^{N} q_i n_i. \quad (1)$$

Here, the number of crossings, $n_i \in [-1, +1]$, was determined by counting ions (with charge $q$) that completely traverse the TMD during the simulation time $\Delta t$. From these counts, the single-channel conductance was calculated as ($G = I_\mathrm{r}/V_\mathrm{e}$), where $V_\mathrm{e}$ is the voltage difference ($V_e = L_z \times$ EF, were $L_z$ is the height of the entire simulation box). The region of the TMD used to calculate ion-crossing events was defined by a right cylinder with a radius of 17.3 Å and a length determined by the z-axis coordinates of the backbone atoms of the two rings of glutamates (positions 258 and 237) at either end of the TMD (Fig. S1).

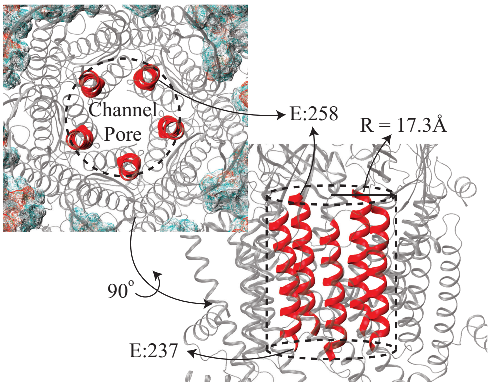


Figure S1: M2 helices lining the TMD pore, having Glu 237 and Glu 258 at the cytosolic and extracellular ends, respectively, are shown as red ribbons. These helices dictate the height of the cylindrical region (with radius of 17.3 Å) used for calculation of the number of ions crossing the TMD.

A $\mathrm{TB_{ion}}$ (top-to-bottom, the subscript denoting the ion type) event occurs whenever an ion crosses the ring of five Glu-258 residues, enters the TMD, and exits through the ring formed by the Glu-237 residues into the ICD; this is the direction of the electric field (orthogonal to the plane of the lipid membrane). Conversely, a $\mathrm{BT_{ion}}$ (bottom-to-top) event occurs whenever an ion moves in the opposite direction, that is, entering from the Glu-237 ring and exiting through the Glu-258 ring. The channel radius was chosen sufficiently wide to capture the obliquity of the cylinder defined by the M2 helices of the five α7 subunits during the simulation and to ensure that an ion never escapes along this height. Given the large conductance of this receptor, only complete events were considered. That is, ions present within the TMD at the start and end of the production phase were disregarded, i.e. $n_i$, is either +1 or −1.

#### S3 Model of Double-Poisson Process

We fitted the distributions of waiting times between events from the simulations to a double-Poisson model. This model comprised of two sequential Poisson distributions, where the output of the first serves as the input of the second distribution. This model assumed events were to be prepared before they conduct subsequently. Starting with the total number of unready events, $N_{\text{unready}}$. Number of events which are ready to conduct, $N_{\text{ready}}$, are define by the lag time before crossing and is governed by the rate of lag, $\lambda_{\text{lag}}$. Events which are ready and conduct are defined by the conduction rate, $\lambda_{\text{cond}}$. The three rate equations of the distributions of these events are

$$\frac{d}{dt}N_{\text{unready}}(t) = -\lambda_{\text{lag}}N_{\text{unready}}, \tag{2a}$$

$$\frac{d}{dt}N_{\text{ready}}(t) = \lambda_{\text{lag}}N_{\text{unready}} - \lambda_{\text{cond}}N_{\text{ready}}, \tag{2b}$$

$$\frac{d}{dt}N_{\text{cond}}(t) = -\lambda_{\text{cond}}N_{\text{ready}}. \tag{2c}$$

A simple integration of the first differential equation gives the distribution of the unready events, $N_{\text{unready}}$, which is an exponential decay with a rate constant $\lambda_{\text{lag}}$. The solution is:

$$N_{\text{unready}}(t) = N_0 \exp\left[-\lambda_{\text{lag}}t\right]. \tag{3}$$

Plugging this expression in the second differential equation, eq. 2b, gives

$$\frac{d}{dt}N_{\text{ready}}(t) = \lambda_{\text{lag}}N_0 \exp\left[-\lambda_{\text{lag}}t\right] - \lambda_{\text{cond}}N_{\text{ready}}.$$

Reshuffling this and multiplying it with the integration factor $\mu = \exp\left[\lambda_{\text{cond}}t\right]$ gives,

$$\exp\left[\lambda_{\text{cond}}t\right]\frac{d}{dt}N_{\text{ready}}(t) - \exp\left[\lambda_{\text{cond}}t\right]\lambda_{\text{cond}}N_{\text{ready}} = \exp\left[\lambda_{\text{cond}}t\right]\lambda_{\text{lag}}N_0\exp\left[-\lambda_{\text{lag}}t\right].$$

Following the Leibniz rule, the expression becomes

$$\frac{d}{dt}\left(\exp\left[\lambda_{\text{cond}}t\right]N_{\text{ready}}(t)\right) = \exp\left[\lambda_{\text{cond}}t\right]\lambda_{\text{lag}}N_0\exp\left[-\lambda_{\text{lag}}t\right].$$

Finally, integrating and simplifying this we get a close form solution for the distribution of number of ready events, $N_{\text{ready}}$:

$$\begin{aligned}
\int \frac{d}{dt}\left(\exp\left[\lambda_{\text{cond}}t\right]N_{\text{ready}}(t)\right)dt &= \lambda_{\text{lag}}N_0\int \exp\left[\left(\lambda_{\text{cond}} - \lambda_{\text{lag}}\right)t\right]dt,\\
\exp\left[\lambda_{\text{cond}}t\right]N_{\text{ready}}(t) &= \frac{\lambda_{\text{lag}}N_0}{\lambda_{\text{cond}} - \lambda_{\text{lag}}}\exp\left[\left(\lambda_{\text{cond}} - \lambda_{\text{lag}}\right)t\right],\\
N_{\text{ready}}(t) &= \frac{\lambda_{\text{lag}}N_0}{\lambda_{\text{cond}} - \lambda_{\text{lag}}}\left(\exp\left[-\lambda_{\text{lag}}t\right] - \exp\left[-\lambda_{\text{cond}}t\right]\right). \qquad (4)
\end{aligned}$$

Using this expression in the last differential equation, eq. 2c, and integrating for the number of conducted events, $N_{\text{cond}}$:

$$\begin{aligned}
\frac{d}{dt}N_{\text{cond}}(t) &= -\frac{\lambda_{\text{cond}}\lambda_{\text{lag}}N_0}{\lambda_{\text{cond}} - \lambda_{\text{lag}}}\left(\exp\left[-\lambda_{\text{lag}}t\right] - \exp\left[-\lambda_{\text{cond}}t\right]\right),\\
N_{\text{cond}}(t) &= -\int \frac{\lambda_{\text{cond}}\lambda_{\text{lag}}N_0}{\lambda_{\text{cond}} - \lambda_{\text{lag}}}\left(\exp\left[-\lambda_{\text{lag}}t\right] - \exp\left[-\lambda_{\text{cond}}t\right]\right)dt.
\end{aligned}$$

The final solution is

$$N_{\mathrm{cond}}(t) = \frac{N_0}{\lambda_{\mathrm{cond}} - \lambda_{\mathrm{lag}}} \left( \lambda_{\mathrm{cond}} \exp\left[-\lambda_{\mathrm{lag}} t\right] - \lambda_{\mathrm{lag}} \exp\left[-\lambda_{\mathrm{cond}} t\right] \right). \tag{5}$$

This gives the distribution of events conducted in time from the total number of events, $N_0$, in the start. Hence, the cumulative distribution function, $F_{\mathrm{events}}(t)$, is

$$F_{\mathrm{events}}(t) = N_0 - \frac{N_0}{\lambda_{\mathrm{cond}} - \lambda_{\mathrm{lag}}} \left( \exp\left[-\lambda_{\mathrm{lag}} t\right] - \exp\left[-\lambda_{\mathrm{cond}} t\right] \right), \tag{6}$$

and the probability density function, $f_{\mathrm{events}}(t)$, is

$$\begin{aligned} f_{\mathrm{events}}(t) &= \frac{d}{dt} F_{\mathrm{events}}(t), \\ &= -\frac{\lambda_{\mathrm{cond}} \lambda_{\mathrm{lag}} N_0}{\lambda_{\mathrm{cond}} - \lambda_{\mathrm{lag}}} \left( \exp\left[-\lambda_{\mathrm{lag}} t\right] - \exp\left[-\lambda_{\mathrm{cond}} t\right] \right). \end{aligned} \tag{7}$$

Number of ready and conducted events were defined to be zero at the start time, $t_i = 0$, and probability is normalized to unity, which implies $N_0 = 1$. Evidently, distributions given in above equations are defined by two independent rate parameters. The time lag before an event occurs is defined by the first parameter, $\lambda_{\mathrm{lag}}$, and rate of conduction is defined by the $\lambda_{\mathrm{cond}}$. These rate parameters of probability density function of the number of conducted events were optimized to the waiting time distribution of the simulated results. Plots of these three distributions with rate parameters as used for the TB and BT waiting distributions from the simulations are shown in Fig. S2.

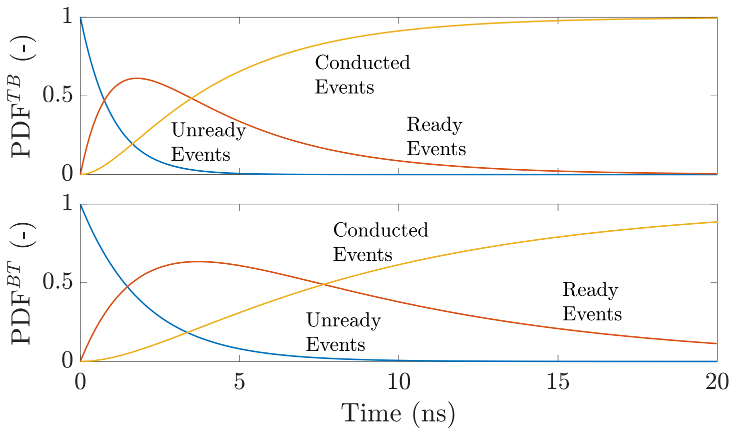


Figure S2: Probability of events over time, at each of the two steps of the double-Poisson model. The top and bottom panels show TB and BT events respectively. The blue plots represent the probability of the total number of unready events (eq. 3). This probability decreases over time as events prepare to conduct after waiting a period, defined as the lag time. The red plots show the probability of events (eq. 4) that have completed the waiting period and are ready to conduct. This distribution increases until events begin to conduct, reflecting the delay introduced by the lag time. Finally, the yellow plots (eq. 5) depict the probability of events that have conducted.

### S4 Single-Channel Conductance: Muscle nAChR

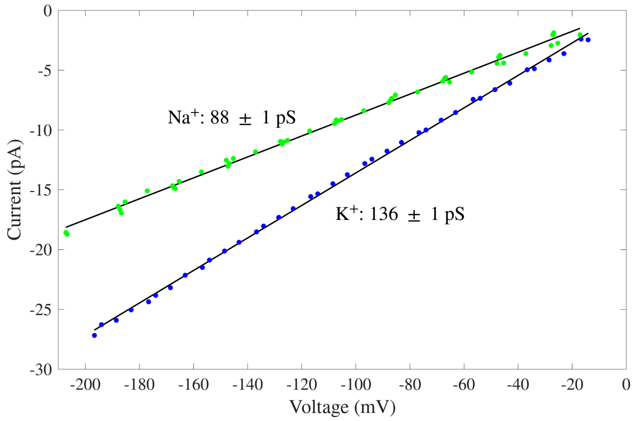


Figure S3: Experimental current–voltage curves recorded from the adult-muscle nAChR. Blue scattered points are $K^+$ inward-current amplitudes recorded from 4 separate cell-attached patch-clamp experiments; green scattered points are $Na^+$ inward-current amplitudes recorded from 5 separate cell-attached patches. The (non-zero) intrinsic membrane potential of each patched cell was accounted for by adding its approximate value to the pipette potential. To increase the accuracy of the current-amplitude estimates, openings of the muscle nAChR were prolonged by engineering the T264P mutation (M2 position 12') in the ε subunit.

#### S5 Channel Conductance: MD Results

*Results included only for PDB IDs with non-zero conductance.*

##### S5.1 Simulations at 22°C

| Index | $TB_{K^+}$ | $BT_{K^+}$ | $TB_{Cl^-}$ | $BT_{Cl^-}$ | Conductance (pS) |
|---|---|---|---|---|---|
| 1 | 31 | 13 | 0 | 1 | 138 |
| 2 | 38 | 13 | 0 | 1 | 189 |
| 3 | 32 | 15 | 0 | 0 | 124 |
| 4 | 42 | 18 | 0 | 0 | 175 |
| 5 | 40 | 6 | 0 | 0 | 248 |
| Average | | | | | |
| - | 37 ± 2 | 13 ± 2 | 0 ± 0 | 0 ± 0 | 175 ± 19 |

Table S9: Summary of events from simulations of 7KOX (unrestrained without ligands) at $V_e = -102$ mV at 22°C. All simulations are 200 ns long. There were no $Cl^-$ ion crossings.

#### S5.2 Restrained Simulations

Restrained MD simulation of each PDB model was 500 ns long, which was broken into sections of five 100 ns to calculate conductance. The results from each run are presented in the table below.

| PBD IDs | $TB_{K+}$ | $BT_{K+}$ | $TB_{Cl-}$ | $BT_{Cl-}$ | Conductance (pS) |
|---|---|---|---|---|---|
| 7EKT | 7 | 0 | 0 | 0 | 102 |
| | 12 | 1 | 0 | 0 | 160 |
| | 10 | 0 | 0 | 0 | 146 |
| | 10 | 0 | 0 | 0 | 146 |
| | 8 | 0 | 0 | 0 | 117 |
| Average | 10 ± 1 | 0 ± 0 | 0 ± 0 | 0 ± 0 | 134 ± 11 |
| 7KOX | 3 | 1 | 0 | 0 | 29 |
| | 1 | 0 | 0 | 0 | 15 |
| | 0 | 0 | 0 | 0 | 0 |
| | 1 | 0 | 0 | 0 | 15 |
| | 0 | 0 | 0 | 0 | 0 |
| Average | 1 ± 1 | 0 ± 0 | 0 ± 0 | 0 ± 0 | 12 ± 5 |
| 8V80 | 1 | 0 | 0 | 0 | 15 |
| | 3 | 0 | 0 | 0 | 44 |
| | 4 | 0 | 0 | 0 | 0 |
| | 3 | 0 | 0 | 0 | 44 |
| | 2 | 1 | 0 | 0 | 15 |
| Average | 3 ± 1 | 0 ± 0 | 0 ± 0 | 0 ± 0 | 24 ± 9 |
| 8V82 | 10 | 2 | 0 | 0 | 117 |
| | 7 | 0 | 0 | 0 | 102 |
| | 12 | 2 | 0 | 0 | 146 |
| | 7 | 0 | 0 | 0 | 102 |
| | 10 | 0 | 0 | 0 | 146 |
| Average | 9 ± 1 | 1 ± 0 | 0 ± 0 | 0 ± 0 | 123 ± 10 |
| 9LH5 | 19 | 14 | 0 | 0 | 73 |
| | 20 | 10 | 0 | 0 | 146 |
| | 17 | 8 | 0 | 0 | 131 |
| | 19 | 11 | 0 | 0 | 117 |
| | 19 | 7 | 0 | 0 | 175 |
| Average | 19 ± 0 | 10 ± 1 | 0 ± 0 | 0 ± 0 | 128 ± 17 |

Table S10: Summary of events and conductance during the restrained PDB structures at $V_e = -102$ mV. Results are from MD simulations ran for 0.5 µs. Each run here is broken into equal sections of 100 ns to calculate conductance.

#### S5.3 Unrestrained Simulations with Ligands

| Index | $TB_{K+}$ | $BT_{K+}$ | Conductance (pS) |
|---|---|---|---|
| 1.a | 39 | 9 | 204 |
| 1.b | 53 | 4 | 333 |
| 2.a | 41 | 10 | 213 |
| 2.b | 57 | 9 | 330 |
| 3.a | 46 | 5 | 281 |
| 3.b | 23 | 2 | 144 |
| 4.a | 46 | 12 | 235 |
| 4.b | 65 | 20 | 310 |
| 5.a | 40 | 9 | 210 |
| 5.b | 61 | 7 | 366 |
| 6 | 19 | 3 | 110 |
| 7 | 35 | 6 | 199 |
| 8 | 49 | 10 | 266 |
| 9 | 10 | 1 | 61 |
| 10 | 43 | 27 | 110 |
| Average | | | |
| - | 41 ± 4 | 9 ± 2 | 225 ± 23 |

Table S11: Summary of events from simulations of 8V82 (unrestrained with ligands) at $V_e = -102$ mV. Longer simulations are broken in intervals of 200 ns (each row is results from 200 ns long simulation). There were no $Cl^-$ ion crossings.

| Index | $TB_{K+}$ | $BT_{K+}$ | Conductance (pS) |
|---|---|---|---|
| 1 | 0 | 0 | 0 |
| 2 | 9 | 1 | 54 |
| 3 | 0 | 0 | 0 |
| 4 | 34 | 0 | 230 |
| 5 | 0 | 0 | 0 |
| Average | | | |
| - | 9 ± 7 | 0 ± 0 | 58 ± 40 |

Table S12: Summary of events from simulations of 8V80 (unrestrained with ligands) at $V_e = -102$ mV. There were no $Cl^-$ ion crossings. Results in each row are from independent 200 ns long simulation.

| Index | $TB_{K^+}$ | $BT_{K^+}$ | $TB_{Cl^-}$ | $BT_{Cl^-}$ | Conductance (pS) |
|---|---|---|---|---|---|
| 1 | 58 | 5 | 0 | 1 | 381 |
| 2 | 60 | 9 | 0 | 2 | 370 |
| 3 | 77 | 12 | 0 | 0 | 455 |
| 4 | 83 | 15 | 0 | 0 | 473 |
| 5 | 73 | 5 | 0 | 0 | 475 |
| Average | | | | | |
| - | 70 ± 5 | 9 ± 2 | 0 ± 0 | 1 ± 0 | 431 ± 21 |

Table S13: Summary of events from simulations of 9LH5 (unrestrained with ligand) at $V_e = -102$ mV.

#### S5.4 Unrestrained Simulations without Ligands

| Index | $TB_{Na^+}$ | $BT_{Na^+}$ | $TB_{Cl^-}$ | $BT_{Cl^-}$ | Conductance (pS) |
|---|---|---|---|---|---|
| 1 | 29 | 14 | 0 | 1 | 113 |
| 2 | 22 | 7 | 0 | 0 | 106 |
| 3 | 30 | 9 | 0 | 0 | 147 |
| 4 | 31 | 9 | 0 | 0 | 153 |
| 5 | 23 | 13 | 0 | 0 | 71 |
| 6 | 18 | 5 | 0 | 0 | 95 |
| 7 | 33 | 7 | 0 | 0 | 189 |
| 8 | 25 | 8 | 0 | 0 | 124 |
| 9 | 33 | 10 | 0 | 0 | 167 |
| 10 | 22 | 6 | 0 | 0 | 117 |
| Average | | | | | |
| - | 27 ± 2 | 9 ± 1 | 0 ± 0 | 0 ± 0 | 128 ± 11 |

Table S14: Summary of events from simulations of 7KOX in NaCl solution (unrestrained without ligands) at $V_e = -102$ mV. Results in each row are from independent 200 ns long simulation.

| Index | $TB_{K^+}$ | $BT_{K^+}$ | Conductance (pS) |
|---|---|---|---|
| 1 | 35 | 0 | 243 |
| 2 | 0 | 0 | 0 |
| 3 | 0 | 0 | 0 |
| 4 | 0 | 0 | 0 |
| 5 | 0 | 0 | 0 |
| Average | | | |
| - | 7 ± 7 | 0 ± 0 | 49 ± 49 |

Table S15: Summary of events from simulations of 8V82 (unrestrained without ligands) at $V_e = -102$ mV. There were no $Cl^-$ ion crossings. Results in each row are from independent 200 ns long simulation.

| Index | $TB_{K^+}$ | $BT_{K^+}$ | $TB_{Cl^-}$ | $BT_{Cl^-}$ | Conductance (pS) |
|---|---|---|---|---|---|
| 1 | 66 | 8 | 0 | 1 | 430 |
| 2 | 46 | 5 | 0 | 0 | 299 |
| 3 | 54 | 13 | 0 | 2 | 313 |
| 4 | 85 | 12 | 0 | 2 | 546 |
| 5 | 72 | 2 | 0 | 0 | 509 |
| Average | | | | | |
| - | 64 ± 7 | 8 ± 2 | 0 ± 0 | 1 ± 0 | 419 ± 44 |

Table S16: Summary of events from simulations of 9LH5 (unrestrained without ligand) at $V_e = -102$ mV.

### S6 Additional Results from MD Simulations

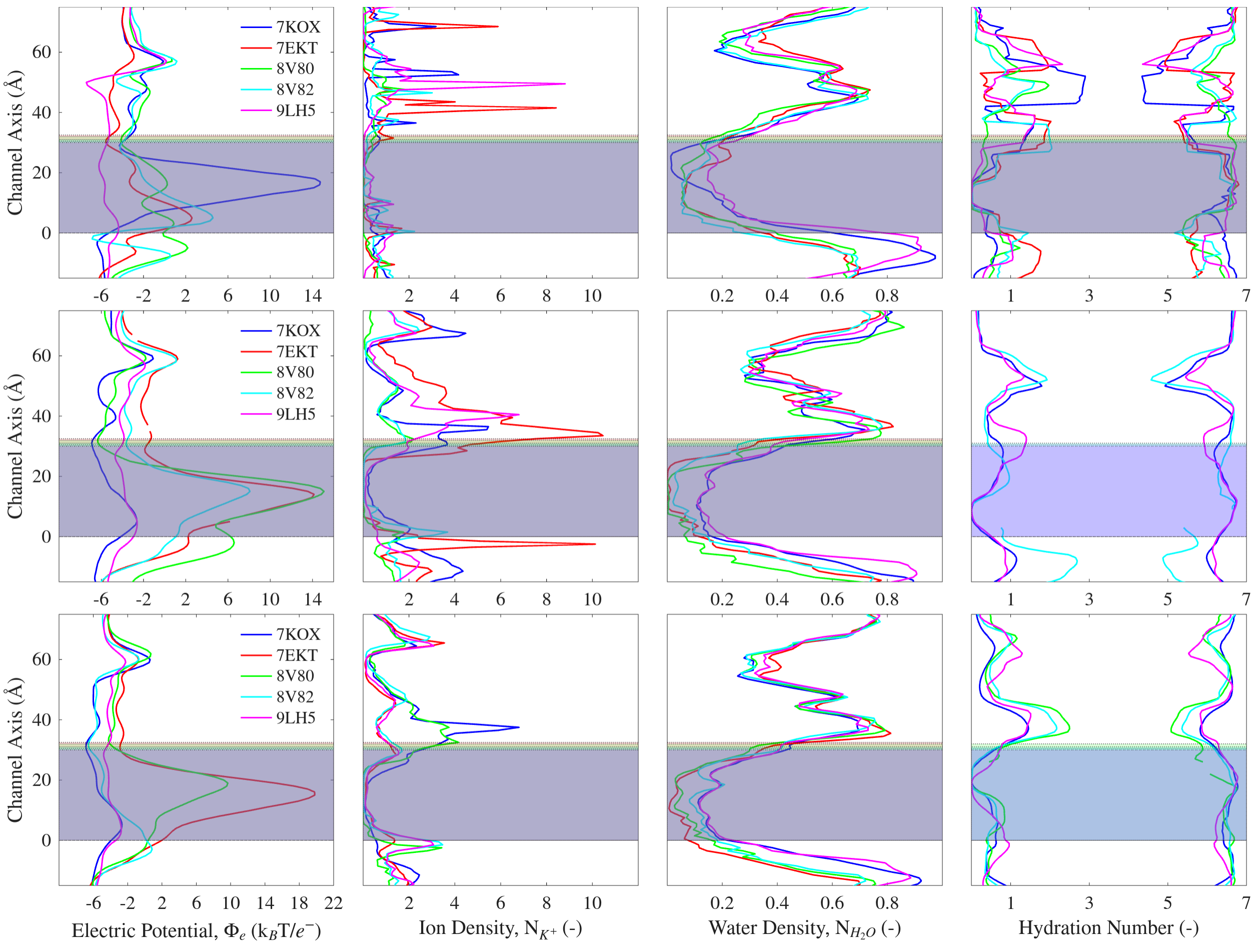


Figure S4: Profiles of the different PDBs structures during the MD simulations, the results in top, middle and bottom row are from simulations when the protein is restrained, unrestrained without ligand, and unrestrained with ligands, respectively. The channel radii of these structures are included in the main text (Fig. 4). The right most plots of ions hydration number is only for structure which have fully or partially hydrated TMD. Water density, in the second last column is the number of water molecules in the channel (identified as a cylinder of radius 17.3 Å) normalized to the number of water molecules found in the same cylinder placed in bulk water.

#### S7 Position Density of Residues

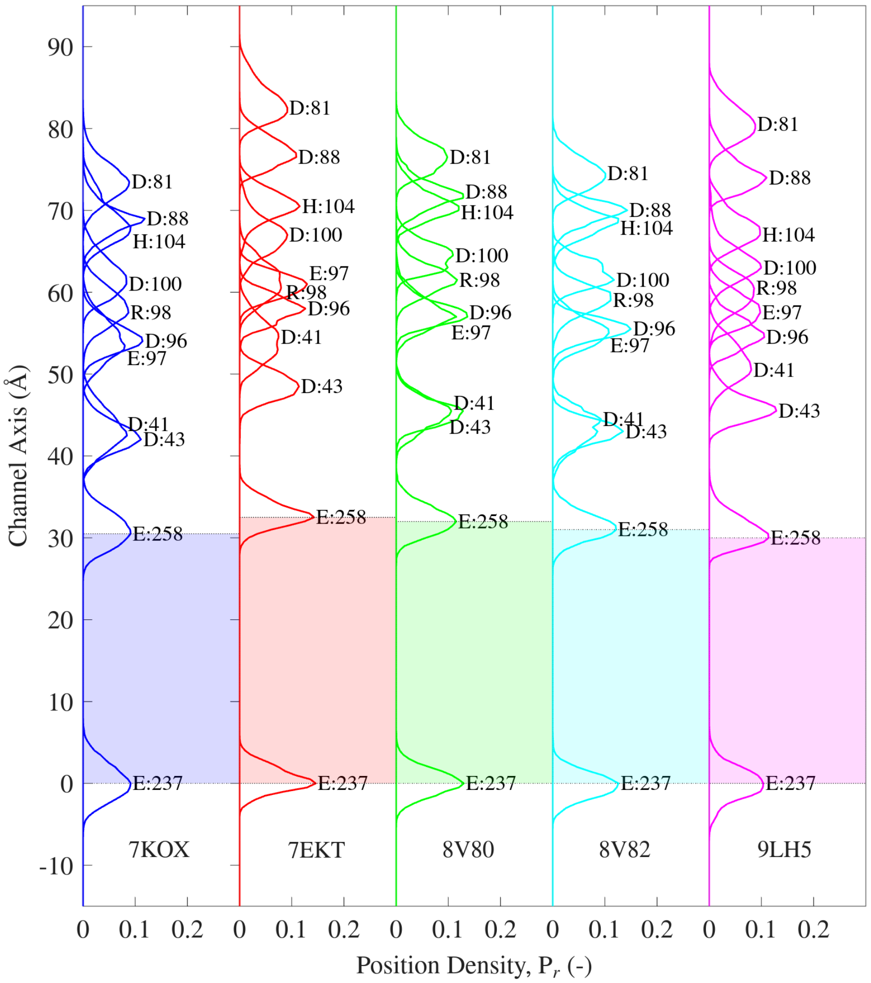


Figure S5: The average position of the charged residues (and His 104) along the pore of the channels from the five simulated structures compared. The shaded regions represent TMDs in each case and structures are vertically referenced to Glu 237 (-1').

#### S8 Variation in Positions of M2 Helix Residues

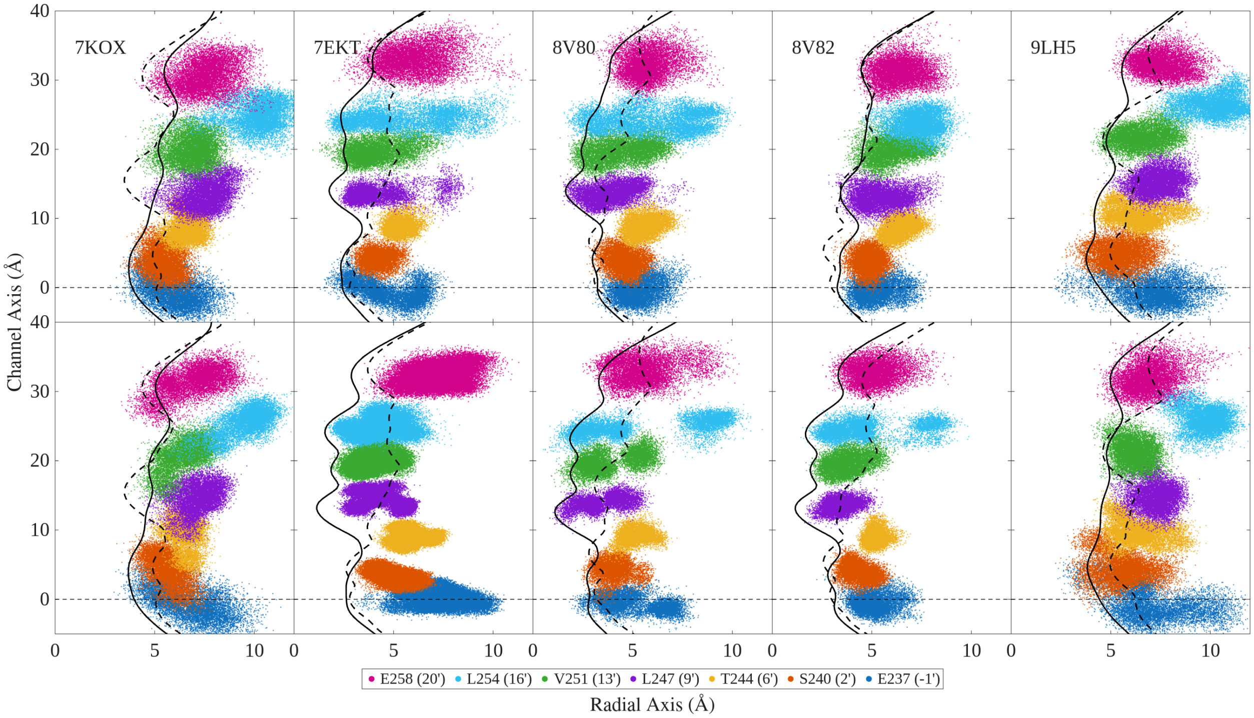


Figure S6: Variations in the side-chains of residues pointing towards the lumen along the M2 α-helix during MD simulation. The top and the bottom row show plots from unrestrained simulation with ligand and without ligands, respectively. Only one simulation, the first run in each case, is chosen to plot data here, except for 8V82 (unrestrained without ligands) when data was plotted for the second run. The first run of 8V82 was conductive, unlike the the rest of the runs for this setup. Each data point is the position of the closest atom to the centre of the pore of each of the 5 subunits. The x-y plane, centred to the middle of the pore, is mapped onto the radial axis and the axis of the channel is referenced to the average position of Glu 237 (-1'). The results shown here are for 200 ns, and positions are plotted every 10 ps. The black solid and dash-dotted curves are the average HOLE profiles from the simulations and the single HOLE profile calculated from the PDB structure, respectively. The two distinct regions of Leu 254 (also two for Val 251 and Leu 247) of PDB 8V80, for example, are due to the ellipticity of the pore, which is most obvious in simulations of this model without ligand. The marked vertical variation is due to the side chains pointing out of the x-y (radial) plane.

### S9 Correlation Analysis between Conductance and Structural Variation

#### S9.1 RMSD Variation between Average Structures

From the 10 individual replicas of the conducting system, PBD ID 7KOX, we constructed the average structures of the receptor using CPPTRAJ using the first 200 ns of each run. The average structure represents the average position of each atom from the simulation run, with reference to the initial structure. The temporal average is calculated every 100 ps. Root mean square displacement (RMSD) of the residues which form the inner lining of the pore between structures from different runs is calculated using MDTraj. This shows how the shape of the average pores varies between structures in different runs. The residues included here are, in order from top of the channel (ECD) to the bottom (ICD), as follows: Lys 8, Lys 12, Val 11, Asn 15, Asp 81, Gln 83, Glu 97, Arg 98, His 104, Lys 45, Asn 46, Asp 43, Glu 258, Glu 237, Gly 236, Ser 240, Thr 244, Val 251.

These RMSD values were sorted by the number of net events for the duration of the simulation and the variation matrix was computed, Fig. S7. If there was a correlation between net events and average structure, then the lowest RMSD would correspond to the smallest difference between net events. However the matrix has a random pattern, showing no obvious relationship between RMSD values of structures and net events.

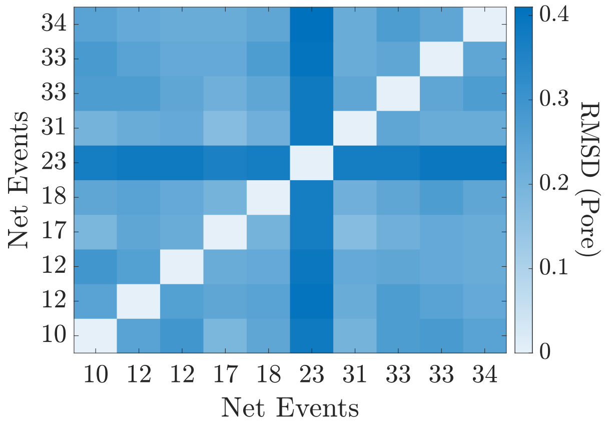


Figure S7: RMSD variation matrix between average structures sorted based on net events from each run at $V_e = -102$ mV.

#### S9.2 PCA and tSNE Analysis

We analyzed the structural states of the α7-nAChR using 120 features of two sets of data, each derived from 10 independent 200 ns simulations (first 200 ns of each replica). These features include pairwise distances between the same residues across all the five chains, with 10 distances per residue. The first data set covers charged residues which face the pore in the ECD and TMDs: Asp 41 (CG atom), Asp 43 (CG atom), Asp 88 (CG atom), Asp 96 (CG atom), Glu 97 (CD atom), Arg 98 (CZ atom), Asp 100 (CG atom), His 104 (CE1 atom), Glu 237 (CD atom), and Glu 258 (CD atom), totalling 100 distance features. We also considered four types of dihedral angles (O-C-CA-CB, C-CA-CB-CG, N-CA-CB-CG, and CA-CB-CG-CD) of Glu 237 on five chains, resulting in additional 20 dihedral angle features in this first data set, composing a total of 120 features. The second data set includes residues within the TMD of the receptor facing the lumen of the channel: Glu 258 (CD atom), Leu 254 (CG atom), Val 251 (CB atom), Leu 247 (CG atom), Val 244 (CB atom), Ser 240 (CB atom) and Glu 237 (CD atom), as shown on Fig. S6, resulting in total of 70 distance features. Before applying machine learning algorithms, we normalized all features by subtracting the mean and dividing by one standard deviation. To reduce dimensionality, we first performed Principal Component Analysis (PCA) using the PCA function from the sklearn.decomposition package. Fig. S8 presents the first and second principal components, with each point representing a state in the configurational space projected onto these components, and each run is coloured according to its net conduction events. No clear correlation was observed between the number of conduction events and configurational states. Since PCA is a linear transformation and may not capture non-linear relationships, we further applied t-distributed stochastic neighbour embedding (tSNE) for visualization in a 2D space, using the TSNE function from the sklearn.manifold package. The TSNE function was applied only to the features from charged residues, first set of data as detailed before. Instead of using the raw 120-feature data, we input the first 50 PCA components into tSNE to mitigate computational costs and account for tSNE's assumption of local linearity, which may not hold in high-dimensional spaces with complex manifolds. The resulting 2D embedding, shown in Fig. S9 and coloured by the number of net conduction events, similarly did not reveal any clear correlation between conduction and configuration. In summary, our PCA and tSNE analyses do not support the hypothesis that variations in conductance arise from differences in the channel's configurational state. Our selected structural features represent only a subset of those necessary to fully characterize the entire protein, yet these selected ones are which directly influence ion crossing through the pore of channel. Expanding the set of features to include more key residues along the conduction pathway may enhance our ability to identify distinct configurational states and further elucidate the relationship between protein structure and conductance.

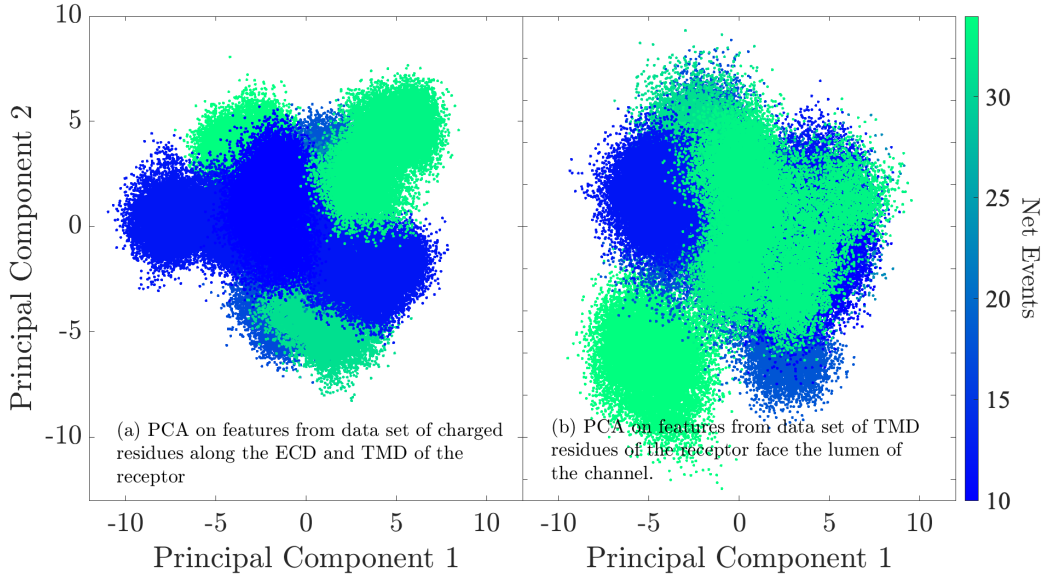


Figure S8: First and second principal components derived from Principal Component Analysis (PCA). Left and right panels show results generated from features of the charged residues along the pore and the TMD facing residues respectively. Each dot represents a snapshot sampled at 10 ps intervals, resulting in a total of 200,000 data points from ten independent simulations. The colour of each dot indicates the net conduction event associated with its respective simulation.

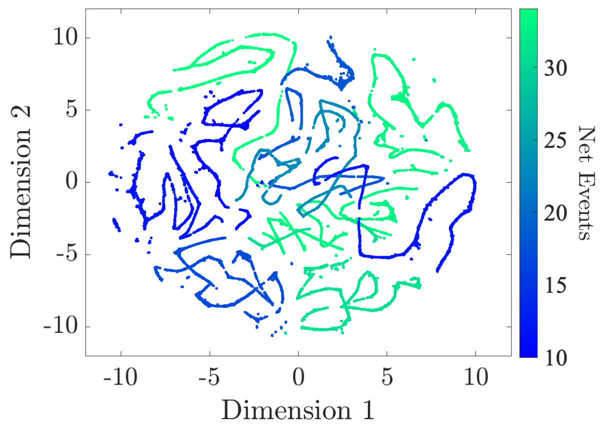


Figure S9: The first and second dimensions derived from t-distributed stochastic neighbour embedding (t-SNE), using the first 50 principal component analysis (PCA) components. Each dot represents a snapshot sampled at 10 ps intervals, resulting in a total of 200,000 data points from ten independent simulations. The colour of each dot indicates the net conduction event associated with its respective simulation.

#### S10 Event Duration

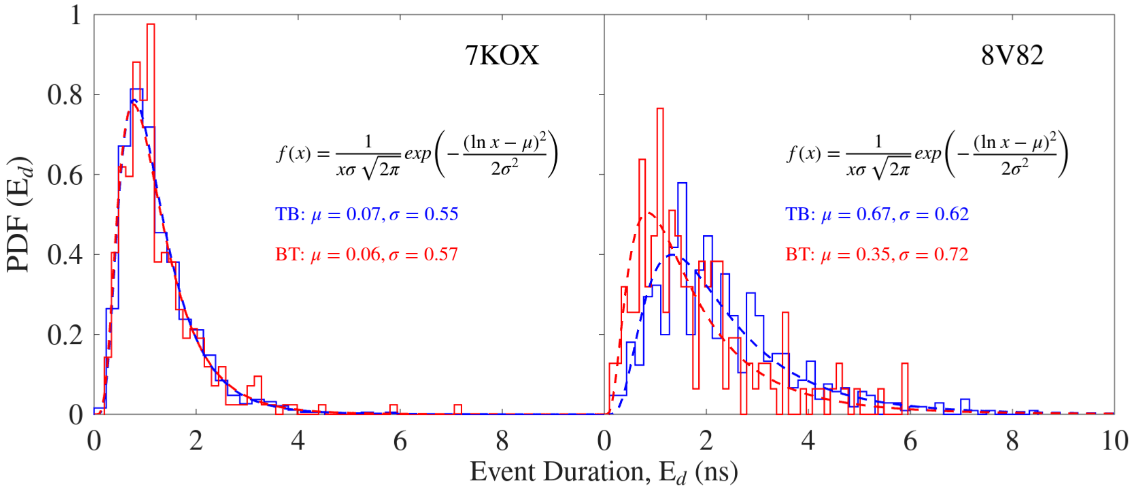


Figure S10: Probability density distribution (PDF) of event duration ($E_d$). This is the duration (in ns) a single $K^+$ ion takes to cross the TMD of the channel, distributions of TB and BT events are colored blue and red respectively. The dashed lines on the plot are log-normal distribution fitted to the data, with location parameters given on the plots. The mean event durations are calculated from the fitted log-normal distribution in each case. The mean duration for 7KOX in both TB and BT directions is 1.24 ns. For 8V82, the duration $K^+$ ions take cross TMD from TB and BT are 2.37 ns and 1.84 ns respectively.

#### S11 Waiting Time for 8V82

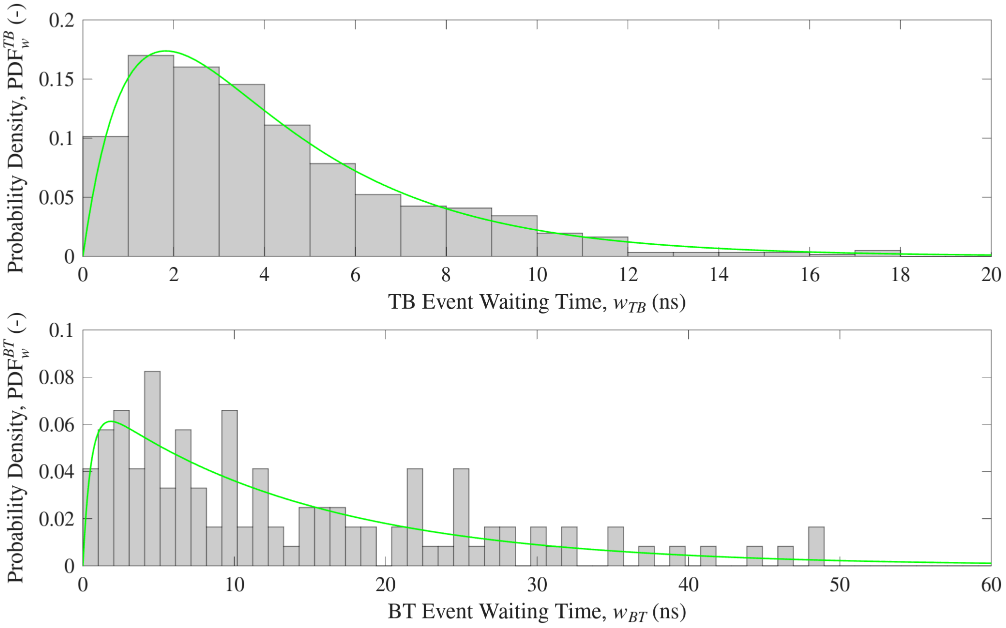


Figure S11: Probability density distributions (PDF) of waiting times between consecutive $TB_{K^+}$ (upper panel) and $BT_{K^+}$ (lower panel) events for unrestrained ligand bound 8V82 (waiting times are calculated from 3 µs worth of simulated data, see table S11). Data fitted in green curves are double-Poisson process, which was hypothesized to best describe the waiting time distribution for this channel. The rate coefficients are $\lambda_{\text{lag}} = 0.9$ and 1.86, and $\lambda_{\text{cond}} = 0.3$ and 0.07 for TB and BT events, respectively.
